# The receptor-like cytoplasmic kinase MAZZA and CLAVATA-family receptors interact *in vivo*, together mediating developmental processes in *Arabidopsis thaliana*

**DOI:** 10.1101/2020.11.12.379859

**Authors:** Patrick Blümke, Jenia Schlegel, Sabine Becher, Karine Pinto, Rüdiger Simon

## Abstract

The receptor-like kinases (RLKs) CLAVATA1 (CLV1) and BARELY ANY MERISTEMs (BAM1 – 3) form the CLV-family (CLVf), which perceives peptides of the CLV3/EMBRYO SURROUNDING REGION (ESR)-related (CLE) family within various signaling pathways of *Arabidopsis thaliana*. CLE peptide signaling, which is required for meristem size control, vascular development, or pathogen responses, involves the formation of receptor complexes at the plasma membrane (PM). These complexes comprise RLKs and co-receptors in varying compositions depending on the signaling context and regulate target gene expression, such as *WUSCHEL* (*WUS*). How the CLE signal is transmitted intracellularly after perception at the PM is not known.

Here, we found that the membrane-associated receptor-like cytoplasmic kinase (RLCK) MAZZA (MAZ) MAZ and additional members of the Pti1-like protein family interact *in vivo* with CLVf receptors. MAZ, which is widely expressed throughout the plant, localizes to the PM via posttranslational palmitoylation potentially enabling stimulus-triggered protein re-localization. We identified a role for a CLV1/MAZ signaling module during stomatal and root development, and redundancy could potentially mask other phenotypes of *maz-1* mutants. We propose that RLCKs such as MAZ mediate CLVf signaling in a variety of developmental contexts, paving the way towards understanding the intracellular processes after CLE peptide perception.

## Introduction

Plant communication and sensing of environmental cues rely on the perception of extracellular signaling molecules via PM-localized cell surface receptors. The genome of *Arabidopsis thaliana* encodes more than 400 membrane-spanning RLKs with intracellular kinase domains (KDs) and versatile types of extracellular domains that perceive e.g. phytohormones, pathogen-associated compounds, or small peptides (Shiu and Bleecker, 2003).

Among this heterogenic group of receptors, CLV1 is one of the best-characterized RLKs with key functions in stem cell regulation in the shoot apical meristem (SAM), inflorescence meristems (IMs), floral meristems (FMs), and the (distal) root meristem (RM) (Stahl and Simon, 2012). Together with its close relatives BAM1, BAM2, and BAM3, CLV1 forms the core family of CLV (CLVf) receptors, which perceive CLE peptides at the PM through their leucine-rich repeat (LRR) receptor domains (Shinohara and Matsubayashi, 2015; Hazak et al., 2017). Beyond meristem homeostasis, CLVf receptors and their associated CLE ligands function in various contexts of plant life, e.g. differentiation of xylem and phloem (Fukuda and Hardtke, 2020), response to bacterial infections (Hanemian et al., 2016), cyst nematode parasitism (Guo et al., 2017), and intercellular spreading of small RNAs or virus particles through plasmodesmata (Rosas-Diaz et al., 2018). Functional diversity of CLE perception via CLVf receptors is also reflected by context-specific interaction and cross-regulation with other RLKs or receptor-like proteins (RLPs).

In the shoot meristems the stem cell derived peptide CLV3, founding member of the CLE family, binds to CLV1 to downregulate the stem cell fate repressing transcription factor WUSCHEL (WUS) as part of a negative feedback loop that dynamically regulates the size of the stem cell domain (Mayer et al., 1998; Schoof et al., 2000; Brand et al., 2000). Besides CLV1, the receptor network that contributes to CLV3 perception in the shoot meristems comprises BAM1 – 3 (DeYoung et al., 2006; Deyoung and Clark, 2008), the heteromeric CLV2/CORYNE (CRN) complex (Jeong et al., 1999; Müller et al., 2008; Bleckmann et al., 2010), and the LRR-RLK RECEPTOR PROTEIN KINASE2 (RPK2) (Kinoshita et al., 2010). BAM1 and BAM2, which were also shown to directly bind CLV3 peptide, can functionally replace CLV1 in the SAM and FM. However, in wild type plants their expression is suppressed in the stem cell domain in a CLV3/CLV1-dependent fashion (Shinohara and Matsubayashi, 2015; Nimchuk et al., 2015). The shoot meristem phenotypes of *clv1* and *clv2/crn* mutants are additive, suggesting synergistic effects in sensing CLV3 (Kayes and Clark, 1998; Müller et al., 2008). Although the CLV2/CRN heteromer resembles the structure of a typical LRR-RLK, the heteromer cannot by itself perceive and transmit CLV3 signals (Nimchuk et al., 2011a; Shinohara and Matsubayashi, 2015).

CLVf receptors are also crucial for CLE perception in the RM. Here, CLV1 is restricted to areas distal to the stem cell organizing quiescent center (QC), where it interacts with the non-LRR-RLK ARABIDOPSIS CRINKLY4 (ACR4) to repress columella stem cell (CSC) fate via a CLE40-dependent pathway (Stahl et al., 2009, 2013). *BAM3* is specifically expressed within the proximal RM in the developing phloem cell files. Here, BAM3 functions as the receptor for CLE45 to regulate sieve element (SE) differentiation together with other phloem-specific factors and the CLV2/CRN heteromer (Anne and Hardtke, 2018; Breda et al., 2019). *BAM1* is broadly expressed in the RM and perceives CLE9/10 peptides to regulate early cell fate decisions during xylem differentiation (Qian et al., 2018).

Co-receptors additionally contribute to the signaling versatility of many LRR-receptor kinase signaling pathways. They are typically characterized by a short extracellular domain with low numbers of LRRs to complete the binding pocket of the corresponding main receptor (recent overview and classification by (Xi et al., 2019)). Mutant analyses and interaction studies revealed that the CLAVATA3 INSENSITIVE RECEPTOR KINASEs1 – 4 (CIK1 – 4) are redundant co-receptors of the CLVf receptors as well as of RPK2 (Hu et al., 2018; Cui et al., 2018; Anne et al., 2018).

The variability of CLVf receptors to function in diverse informational networks with distinct signaling outputs requires mechanisms that add specificity to each pathway. One potential layer of specificity can arise from downstream signaling. However, the immediate early events after recognition of CLE peptides by CLVf receptors are not clear. Studies in maize and Arabidopsis suggest the involvement of heterotrimeric GTP binding protein (G-protein) complexes to integrate CLE peptide responses (Bommert et al., 2013; Ishida et al., 2014; Wu et al., 2020). Mitogen-activated protein kinase (MAPK) cascades, which display a classical mode of intracellular signal transduction in RLK pathways (reviewed in (He et al., 2018)), have been proposed to mediate downstream CLE signaling (Betsuyaku et al., 2011; Lee et al., 2019). However, no physical interaction between CLVf receptors and MAPK signaling elements have yet been shown.

Many cell surface receptor-based signaling pathways rely on RLCKs as key mediators of information transduction, integration, and attenuation (reviewed in (Liang and Zhou, 2018)). RLCKs form a heterogenous group of signaling kinases that are characterized by the absence of extracellular receptor domains and lack a TMD in most cases, however, they can be PM-associated (Bi et al., 2018). The Arabidopsis genome contains 402 potential *RLCK*-encoding genes that cluster into 15 phylogenic sub-groups (Shiu et al., 2004; Fan et al., 2018). These comprise proteins like the RLCK class VII member BOTRYTIS-INDUCED KINASE1 (BIK1), which acts as an important signaling hub downstream of various LRR-RLKs to control immune responses, e.g. via FLAGELLIN SENSING2 (FLS2), but also mediates processes of developmental regulation through BRASSINOSTEROID INSENSITIVE1 (BRI1) or the ERECTA-family (ERf) (Lu et al., 2010; Lin et al., 2013; Chen et al., 2019). Moreover, non-LRR-RLKs also depend on RLCKs. For example. FERONIA (FER), ANXUR1 (ANX1), or ANX2 from the *Catharanthus roseus* RLK1-like (CrRLK1L) family regulate cell wall (CW) integrity through MARIS (MRI) (Boisson-Dernier et al., 2015). MRI belongs to the RLCK VIII subgroup, which contains 11 Arabidopsis genes that share high sequence similarity with *Pti1* from *Solanum lycopersicum* (tomato), a target of the Pto kinase conferring resistance against the bacterial speck disease (Zhou et al., 1995; Sessa et al., 2000). Furthermore, members of the Pti1-like proteins in Arabidopsis (Pti1-1, Pti1-2, Pti1-3, Pti1-4) were shown to interact with OXIDATIVE STRESS INDUCIBLE1 (OXI1), which mediates the cellular response to stress signals like ROS and fungal elicitors. Pti1-4 was also associated with MAPK signaling downstream of OXI1 (Anthony et al., 2006; Forzani et al., 2011). Additionally, the Pti1-like family members CYTOSOLIC ABA RECEPTOR KINASE1 (CARK1) and CARK6 were found to be interactors of the REGULATORY COMPONENTS OF ABA RECEPTORS (RCARs) that function in perceiving the phytohormone abscisic acid (ABA), a general trigger of abiotic stress responses (Zhang et al., 2018; Wang et al., 2019).

Although some members from the RLCK VIII family in Arabidopsis have emerged as crucial signaling intermediates of several pathways, most Pti1-like homologs are not characterized in detail, or not at all. We here report the identification of MAZZA (MAZ, Pti1-3), a member of the RLCK subgroup VIII, as an interactor of the CLVf receptors, thereby expanding the potential range of the RLCK VIII / Pti1-like family to LRR-RLK pathways. We found that CLV1 and MAZ together are involved in developmental processes within stem cell regulation in the RM and during stomatal patterning.

## Results

### MAZ interacts with CLV1 *in vivo*

CLVf receptors are critical for meristem homeostasis but also play pivotal roles during other developmental and physiological signaling events. This is, for example, reflected in the expression of *CLV1* in various tissues besides the shoot meristems, like companion cells (CCs) of the phloem in aerial organs and the root, and in the distal RM. Furthermore, we found that *CLV1* is expressed in the epidermis of cotyledons and true leaves, particularly in developing cells of the stomata lineage (Fig. 1, Suppl. Fig. 1 A). However, direct downstream targets of CLVf receptors are not known in any of those tissues. To identify CLV1 interactors we applied an untargeted, non-organ-biased co-immunoprecipitation (CoIP) approach with magnetic anti-GFP beads (ChromoTek) against CLV1-2xGFP from whole Arabidopsis seedlings expressing the receptor-fluorophore fusion under the control of the endogenous *CLV1* promotor. This transgene is fully functional and was previously shown to complement the known phenotypes of *clv1* mutants (Nimchuk et al., 2011b). Via subsequent mass spectroscopy analyses of the Co-IP fractions we detected Pti1-like proteins among the peptide sequences specifically pulled down with CLV1 and not found in GFP controls (Suppl. Tab. 1). Most of those hits were unique for Pti1-3, while some identified peptide sequences also aligned to other Pti1-like members. Among the candidates for CLV1 interaction we identified Pti1-3 as most promising due to its predicted PM-localization and kinase function. The *Pti1-3* gene (At3g59350) encodes a protein with a predicted KD and an undefined N-terminal region. Due to its capability to interact with CLV1 *in vivo* we dubbed the protein MAZZA (MAZ, Italian for “club”, in analogy to the Latin “clavata”, and Greek “coryne”).

**Figure 1.**
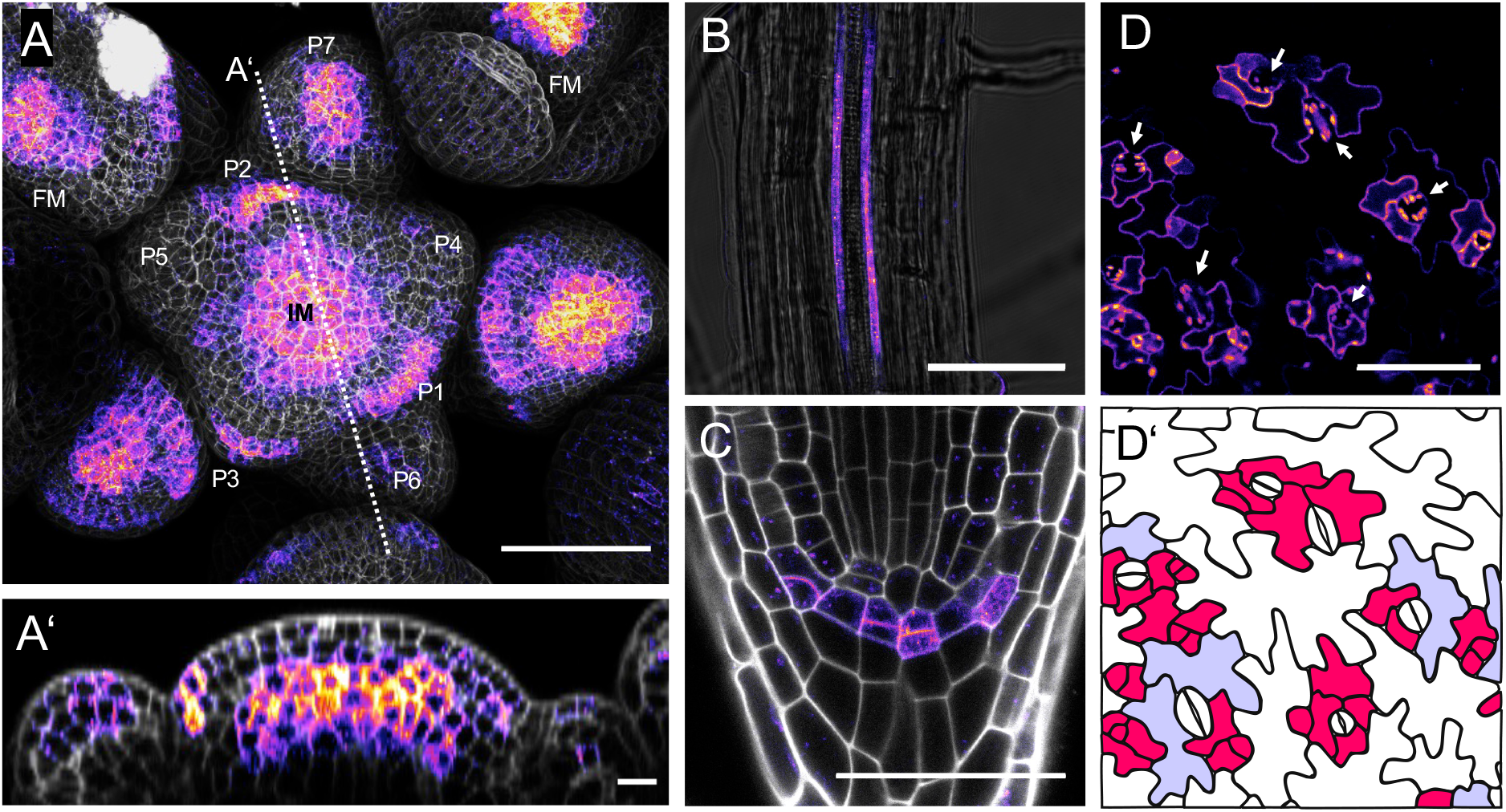
Expression patterns of *CLV1:CLV1-2xGFP* in different tissues of Arabidopsis. **A, A’** Inflorescence with inflorescence meristem (IM), floral primordia, and floral meristems (FM) of 5 weeks old *CLV1*:*CLV1-2xGFP//clv1-11* plants depicted as maximum intensity projection of a z-stack (A) and XZ section (A’) at indicated position. *CLV1* is expressed in the central zone of the meristems and in border cells toward newly arising primordia. **B** Root section within the differentiation zone of a seedling 5 DAG shows presence of the fusion protein CLV1-2xGFP in companion cell files (compare Suppl. Fig. 1 A). **C** Within the root meristem *CLV1* expression is restricted to columella stem cells and epidermis/lateral root cap initials (seedling 5 DAG). **D** In the epidermis of first true leaves (14 DAG) *CLV1-2xGFP* expression is predominantly found in cells of the stomata lineage. Guard cells (white arrows in D) show no expression at the PM (note autofluorescence of chloroplasts). **D’** The schematic representation of D illustrates outlines of all cells (based on bright field channel). Cells with clear GFP fluorescence signals at the PM are colored in pink, with weak signals in light blue. A, A’, C: merge of GFP channel (LUT: fire) and PI (grey). B: merge of GFP (fire) with bright field (grey). Scale bares: 50 μm (A, B, C, D) and 10 μm (A’).

MAZ is also found in complex with CLV1 when following a targeted CoIP strategy with leaf material from *Nicotiana benthamiana* plants, transiently expressing the respective interaction partners fused either to GFP or mCherry (mCh) to allow subsequent immunodetection (Fig 2 A). Supplying CLV3 peptide prior to protein extraction did not influence complex formation with CLV1, pointing towards a constitutive, ligand independent interaction. To analyze the complex at subcellular level, we applied Förster resonance energy transfer (FRET)-based fluorescence-lifetime imaging microscopy (FLIM) and demonstrated direct interaction between MAZ and CLV1 at the PM (Fig. 2 B). In these FLIM experiments within transiently transformed *N. benthamiana* leaf epidermis cells, the donor-lifetime of CLV1-GFP decreases significantly in the presence of MAZ-mCh (mean FRET efficiency of 5.9 %, Supp. Tab. 2), while a PM-located negative control tagged to mCh affects donor lifetime only at a mean FRET efficiency of 2.0 %, which we defined as the background level for randomly distributed acceptor fluorophores in all following FLIM assays. Infiltration of 5 μM CLV3 peptide into the leaves before FLIM measurements did not affect lifetime. This suggests that CLV3 ligand binding to CLV1 neither enhances the interaction with MAZ, nor induces a rapid dissociation of the two proteins within this experimental setup.

**Figure 2.**
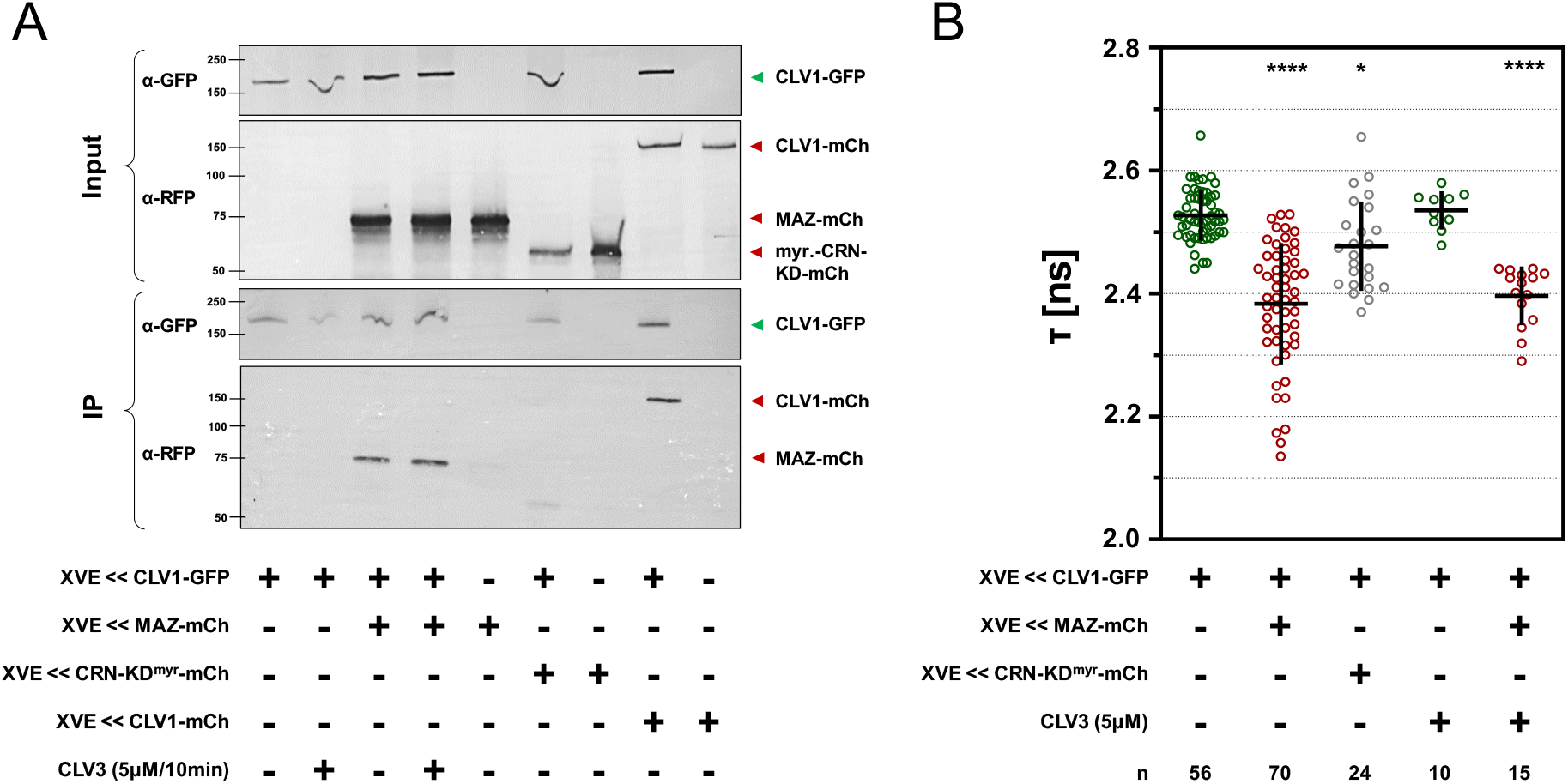
*In vivo* interaction analysis of MAZ and CLV1 via CoIP and FLIM. CoIP and FLIM was performed with *N. benthamiana* leaf epidermis material transiently expressing combinations of indicated fluorophore fusions under the control of the *XVE<<oLexA-35S* estradiol inducible system. **A** MAZ-mCherry (mCh) is co-immunoprecipitated with CLV1-GFP using anti-GFP magnetic beads and subsequent immunodetection with antibodies against GFP and RFP, respectively. The negative control, myristoylated CRN-KD-mCh, is not pulled down together with CLV1-GFP, while the detection of CLV1-mCh reflects the previously reported homomerization of CLV1 (Bleckmann et al., 2010). Treatment of CLV3 peptide (5μM, 10 min prior to sample preparation) does not affect the interaction in this setup. **B** The fluorescence lifetime τ [ns] of CLV1-GFP is reduced in the presence of MAZ-mCh, while the PM control (CRN-KD^myr^-mCh) reduces the τ of CLV1-GFP with low efficiency. Infiltration of a 5 μM CLV3 peptide solution into the leaves 5 – 15 min prior to measurements does not alter the τ values of donor-only nor FRET samples. Number of repetitions are indicated, p-values were calculated by ANOVA and Dunnett’s post hoc test, with * for p ≤ 0.05, and **** for p ≤ 0.0001. Error bars display SD.

### MAZ and Pti1-like homologs interact with CLVf receptors and related pathway elements

Next, we extended the FRET-FLIM interaction studies in the *N. benthamiana* expression system to test if MAZ can interact with other proteins that contribute to CLE peptide perception. We detected interactions at the PM between MAZ and the CLVf receptors BAM1 and BAM3 with mean FRET efficiencies of 6.5 % and 4.0 %, respectively (Fig 3 A, B, Suppl. Tab. 2). Furthermore, MAZ interacts with the co-receptor CIK2 (5.7 %, Fig 3 A), and with CRN in the presence of CLV2 (3.0 %, Fig. 3 B). These data indicate that MAZ could integrate different CLE peptide triggered RLK pathways.

**Figure 3.**
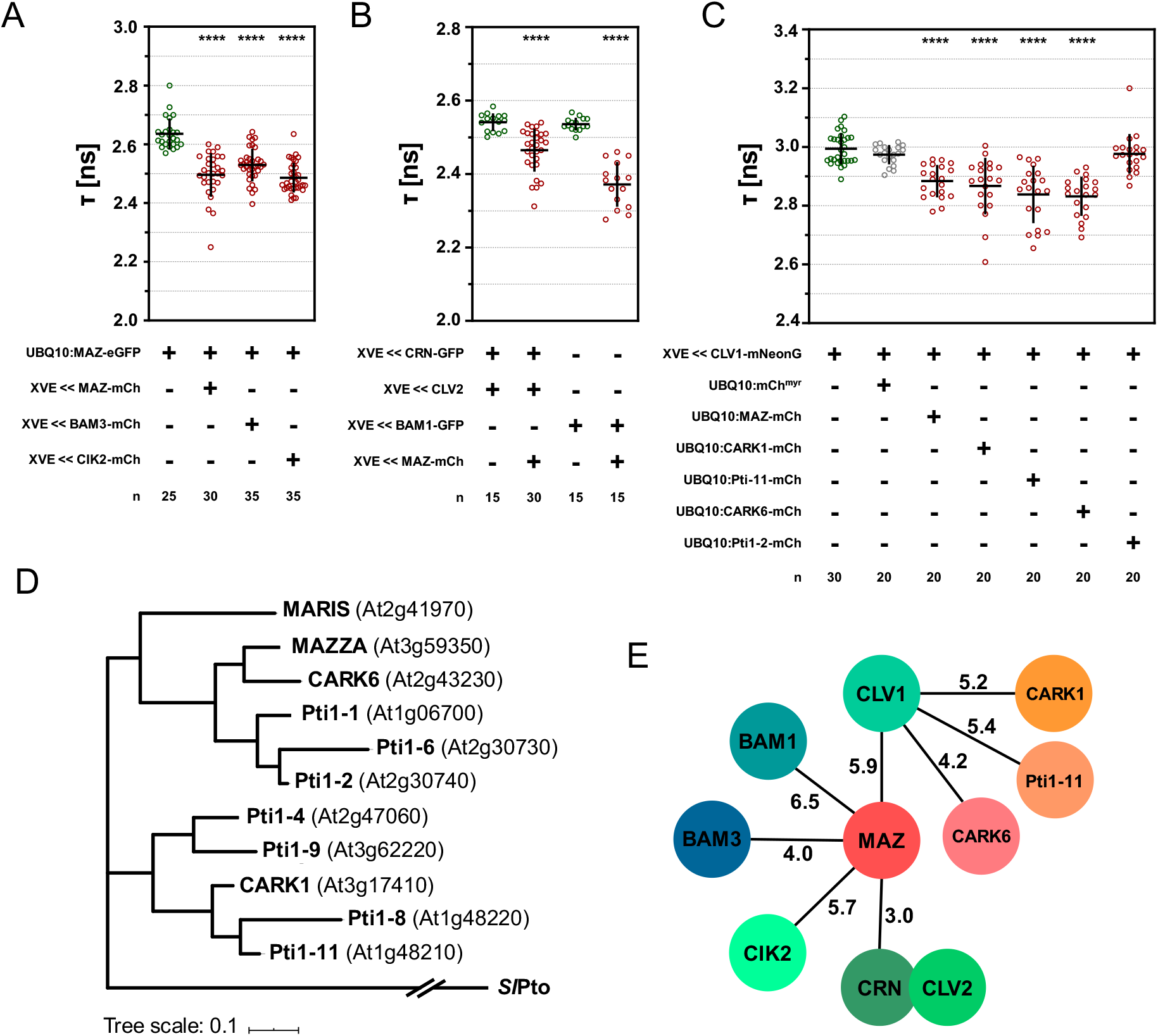
FLIM interactome of CLVf receptors and related signaling elements with the Pti1-like family. **A – C** Fluorescence lifetime τ [ns] was determined in the *N. benthamiana* system, expressing the indicated constructs via the constitutive *UBQ10* promoter or under the control of the *XVE<<oLexA-35S* estradiol inducible module. **A** Donor τ of MAZ-eGFP is reduced in the presence of MAZ-mCh, BAM3-mCh, and CIK2-mCh, respectively. Note that the enhanced GFP variant (eGFP) was used here, explaining the slight differences compared to donor-only τ values in Fig 2 B and 3 B with constructs harboring the regular (F64) GFP. **B** The τ of CRN-GFP is decreased significantly but with a comparable low mean FRET efficiency of 3 % in the presence of MAZ-mCh and CLV2, while τ of BAM1-GFP is reduced with a FRET efficiency of 6.5 % by MAZ-mCh. **C** The τ of CLV1-mNeonGreen is significantly reduced in the presence of MAZ-mCh, CARK1-mCh, Pti1-11-mCh, CARK6-mCh, but not in combination with Pti1-2-mCh (nor by the negative control myristoylated mCh). Number of repetitions for FLIM experiments are indicated, p-values were calculated by ANOVA and Dunnett’s post hoc test, with **** for p ≤ 0.0001. Error bars display SD. **D** Phylogram of the Pti1-homologs of *A. thaliana* and the *Solanum lycopersicum* SlPto as outgroup. The tree was generated after initial multiple sequence alignment (ClustalΩ) and subsequently applying a Maximum Likelihood strategy in MEGAX (via the JTT-matrix based model and 1000 repetitions). Final tree visualization was realized with iTOL software. Branch length is in scale as indicated displaying the calculated substitutions/site rate. **E** Graphical summary of the FLIM interactions identified here, mean FRET efficiency values are indicated for each pair (compare Suppl. Tab. 2).

Additionally, lifetime of MAZ-eGFP decreases in the presence of MAZ-mCh with a mean FRET efficiency of 5.3 % (Fig.3 A). Thus, MAZ can form dimers or higher ordered homomers, which might reflect auto-regulative mechanisms.

Since all 11 members of the Pti1-like family in Arabidopsis are highly conserved at amino acid (aa) sequence level (Appendix), they might be functionally redundant. Indeed, from four Pti1-like homologs that we tested additionally to MAZ, three, namely CARK1, CARK6, and Pti1-11 were able to interact with CLV1 at the PM of *N. benthamiana* leaf epidermis cells with mean FRET efficiencies ranging from 4.2 to 5.4 %. Only Pti1-2 showed no interaction with CLV1 (Fig. 3 C). This observation is not reflected in the phylogeny of the Pti1-like family, because, for example, CARK1 is more distantly related to MAZ than Pti1-2 (Fig. 3 D). However, differences in the phylogeny do not necessarily concern the sequences that are crucial for protein complex formation. *Vice versa*, detecting no change of donor lifetime in the presence of a certain acceptor does not generally exclude physical interaction, e.g. if the fluorophore fusion sterically hinders energy transfer.

### Subcellular localization of MAZ at the PM depends on palmitoylation

MAZ interacts with RLKs and other signaling proteins at the PM. Accordingly, all analyzed MAZ fusion proteins with different fluorophores are PM-localized after transgene expression in *N. benthamiana* (Suppl. Fig. 3, 4) as well as in *A. thaliana* (Fig. 4, 5 Suppl. Fig. 1, 6, 7). MAZ co-localization with CLV1 at the PM is not extended to CLV1-GFP vesicles in *N. benthamiana* cells, which indicate receptor sequestration (Suppl. Fig. 3). This suggests that MAZ is not sequestered together with CLV1 but might instead follow distinct signaling routes after RLK activation and turnover.

**Figure 4.**
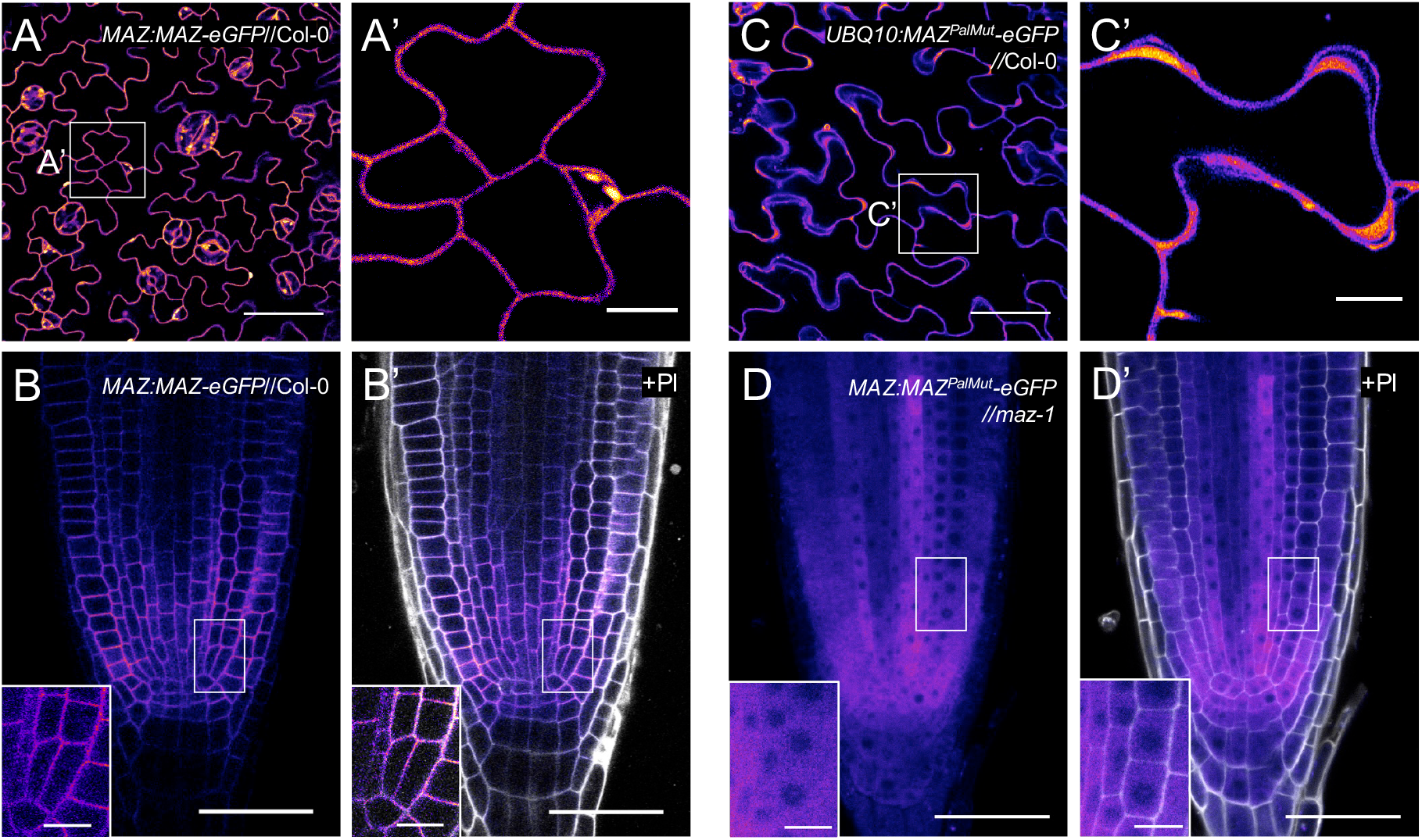
Subcellular localization of MAZ at the PM depends on conserved palmitoylation sites. Stable expression of different MAZ variants demonstrate the impact of the predicted palmitoylation sites C48, C49 (and the adjacent C51) on subcellular localization of MAZ-eGFP. **A – B’** The wild typic *MAZ* protein (fused to eGFP) is located at the PM after expression under the native *MAZ* promotor, as shown here in rosette leaves (A, A’) and in the root meristem (RM, B, B’). **C – D’** Site directed mutagenesis of the palmitoylation sites causes a shift of localization of the mutant MAZ^PalMut^-eGFP fusion protein from the PM to the cytoplasm, both, when expressed under the *UBQ10* promotor, e.g. in leaf epidermis cells (C, C’’), or under the control of the native *MAZ* promotor, shown here in the RM region. Sale bars: 50 μm (A, B, B’, C, D, D’), 10 μm (A’, C’, detail views of B, B’, D, D’). All images are displayed in the LUT fire, in B’ and D’ the merge with PI (grey) is shown.

**Figure 5.**
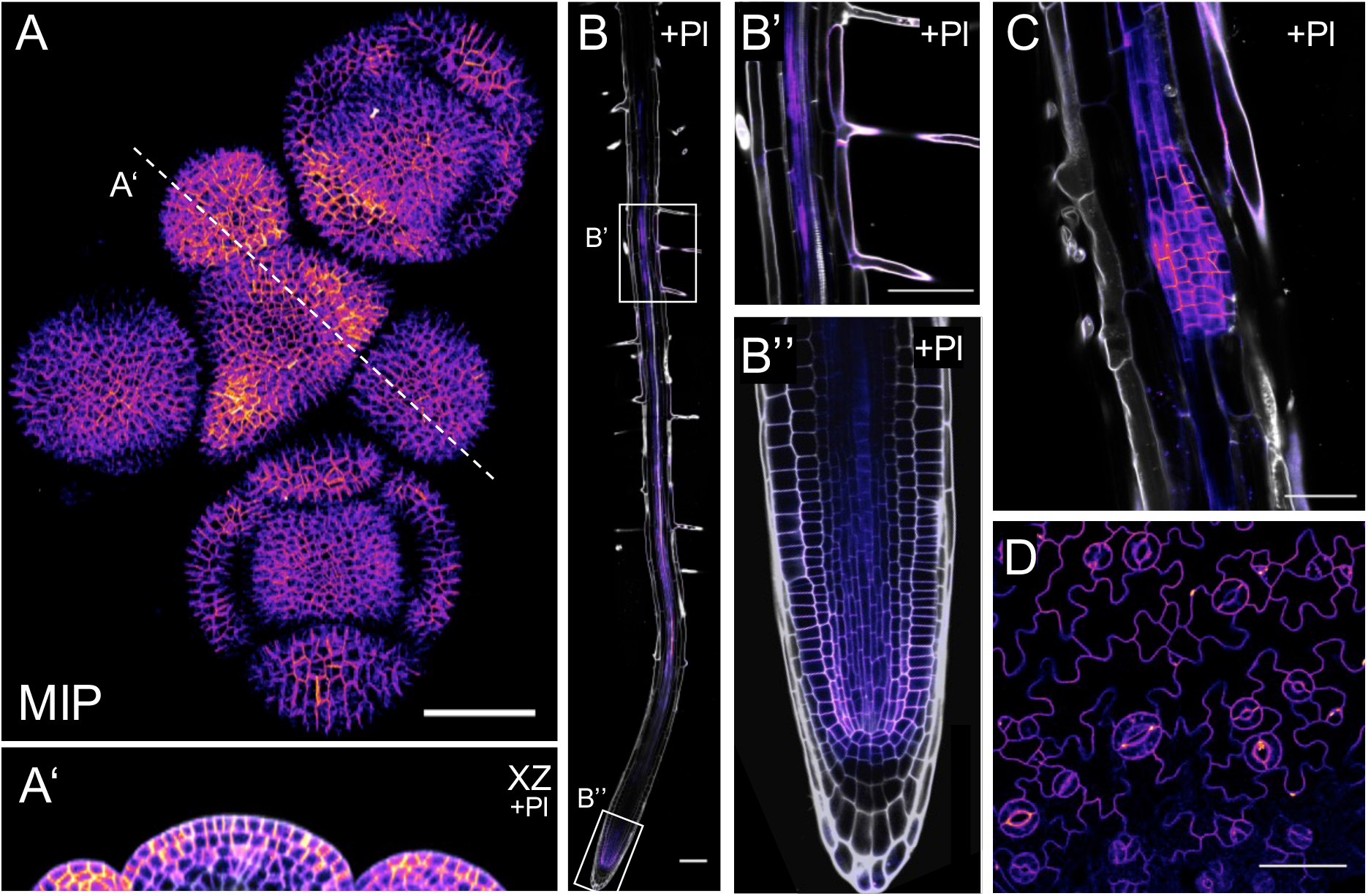
Expression analysis of *MAZ:MAZ-eGFP*//Col-0. **A, A’** Within the inflorescence, MAZ-eGFP is found in all cell layers. Z-stack acquired from a sample of a 5 weeks old plant, presented here as maximum intensity projection ( MIP, A) and XZ section at indicated position (A’). *MAZ* expression is elevated at boundary regions towards newly formed primordia and towards organs in later stages of development. **B, B’, B’’***MAZ* expression in the root (imaged 5 DAG) comprises the vasculature, root hairs (B’), and the root meristem (B’’). **C** Emerging lateral roots show distinct fluorescence signals of the MAZ-eGFP fusion (10 DAG, compare Suppl. Fig. 6). **D** In the epidermis of cotyledons (14 DAG), MAZ-eGFP is present in pavement cells, in the stomata lineage, and in mature guard cells. Scale bars: 50 μm (A, B’ – D), 10 μm (A’), 100 μm (B). GFP-signals visualized by the LUT fire, in A’ – C merge with PI (grey).

**Figure 6.**
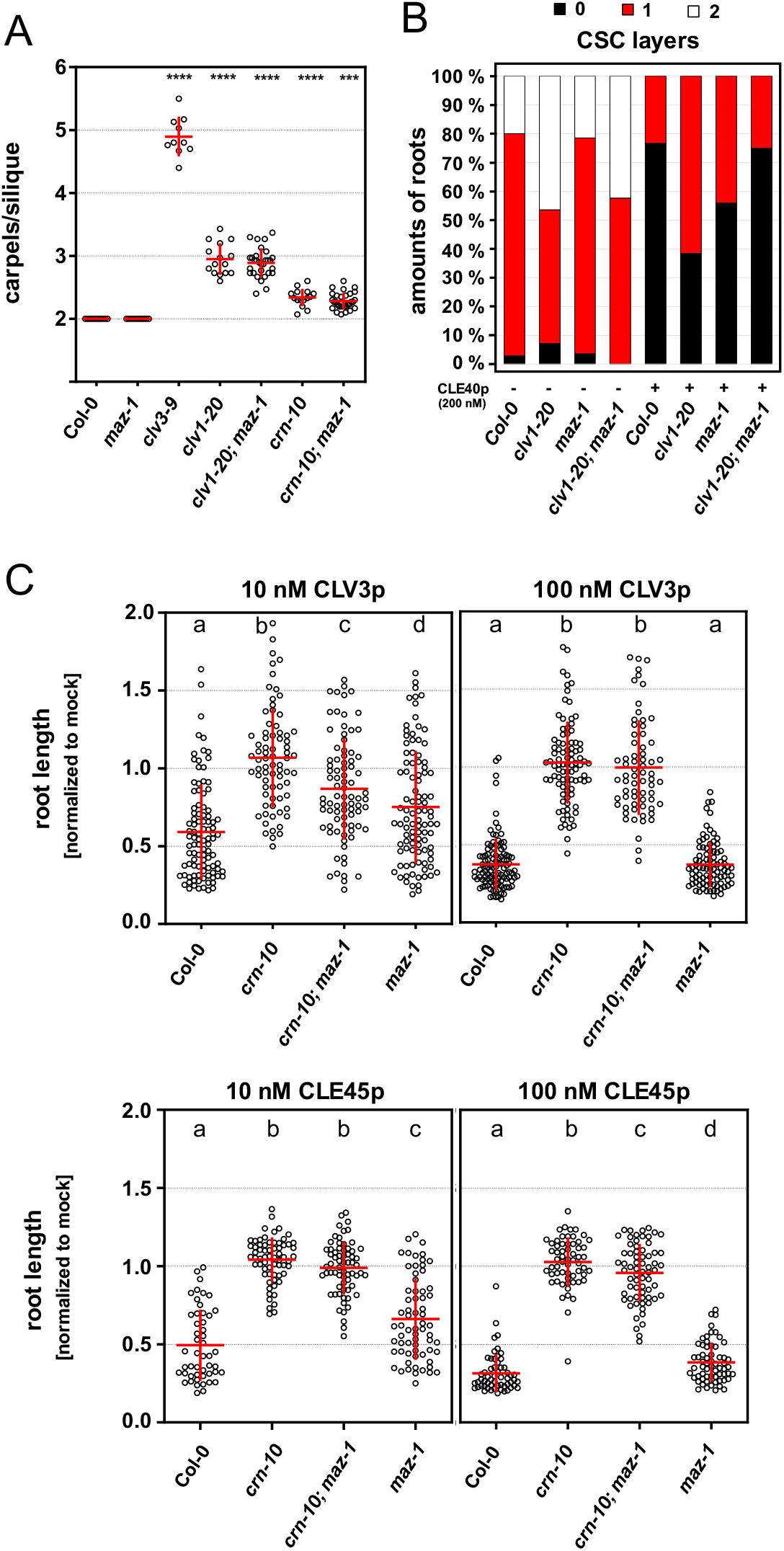
*Maz-1* mutants show no increased carpel number but are partially resistant against CLE peptide treatment. **A** Mean carpel number per silique of indicated genotypes monitored in 6 – 8 weeks old plants grown under long day conditions. For each genotype 10 – 30 plants were analyzed by counting the carpels of 15 – 30 siliques per plant. The average carpel numbers for each plant are plotted. P-values calculated by ANOVA and Dunnett’s post hoc test with *** for p≤ 0.001, and **** for p ≤ 0.0001. Sample mean and SD displayed in red. **B** CSC layers of indicated mutants quantified after mPS-PI staining of seedlings 5 DAG grown on ½ MS agar plates with and without 200 nM CLE40 peptide. **C** Root length of different genotypes 10 DAG grown on ½ MS agar plates supplemented with CLV3p or CLE45p at 10 nM and 100 nM, respectively. Data are normalized to the mean of the corresponding genotype grown without peptide. Statistical groups were assigned after calculating p-values by ANOVA and Dunnett’s post hoc test (differential grouping from p ≤ 0.05). Sample mean and SD displayed.

**Figure 7.**
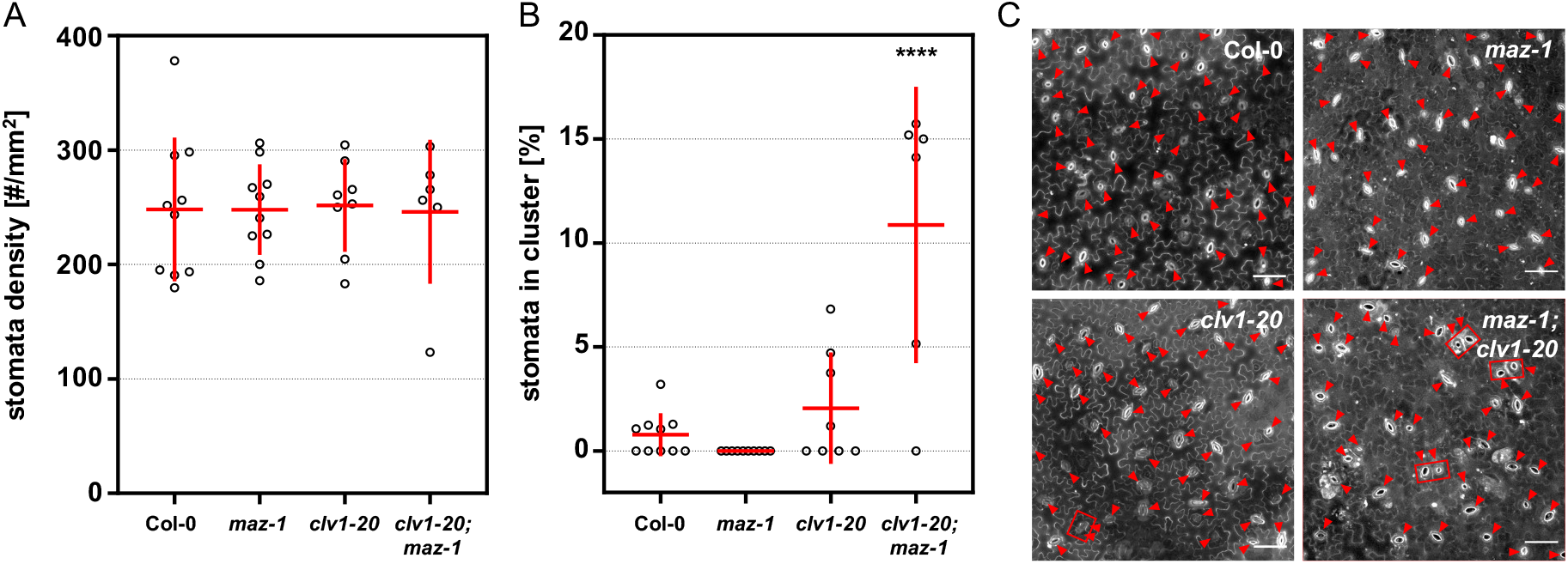
*Clv1-20;maz-1* double mutants display significantly increased stomata cluster rates. **A** Stomatal density and **B** cluster rate were accessed from true leaves of indicated genotypes. Each plotted data point represents the mean value of 4 areas within one leaf of individual plants. Seedlings were grown on ½ MS agar plates. 14 DAG leaves were stained with PI to image abaxial epidermis cells. Significant differences in comparison to the Col-0 sample are indicated with **** for p ≤ 0.0001, calculated by ANOVA and Dunnett’s post hoc test. Sample mean and SD displayed with red lines. **C** Representative images of the analyzed leaf epidermis areas counterstained with PI. Arrow heads mark stomata and boxes indicate clusters. Scale bars: 50 μm.

The MAZ protein sequence does not harbor any TMD, but the N-terminal region possesses predicted palmitoylation sites (C48, C49, Suppl. Tab. 3). S-acylation, i.e. covalent binding of palmitic acid or other fatty acids to the thiol group of cysteine residues, is reversible and thus can represent a mechanism of specific regulation (Hurst and Hemsley, 2015). We found that the predicted MAZ palmitoylation sites are responsible for its PM-localization. Exchanging the corresponding cysteines in the fusion construct *MAZ*^*PalMut*^-*eGFP* results in high amounts of cytoplasmic fluorescence signals instead of PM-localization of the fusion protein after expression in *N. benthamiana* or Arabidopsis (Fig 4, Suppl. Fig. 4). This strongly indicates that subcellular PM localization of MAZ relies on post-translational modification via S-acylation.

### *MAZ* shares expression domains with the CLVf receptors in *A. thaliana*

*MAZ* is expressed in various tissues and organs of Arabidopsis including the distinct expression domains of the CLVf receptors. Analyzing several independent fluorescent *MAZ* reporter lines, we observed differences between translational and transcriptional reporters suggesting that the *MAZ* coding sequence may affect protein expression (Suppl. Fig. 5). Thus, in the following experiments we used translational lines comprising the *MAZ* promotor sequence and the *MAZ* coding sequence including introns to study *MAZ* expression.

In the shoot, *MAZ* is expressed in cells of the leaf epidermis, including stomata precursor cells and guard cells (GCs), the vasculature, and the hypocotyl epidermis (Fig 5 D, Suppl. Fig. 1, 6 C). Furthermore, within the inflorescence we detected *MAZ* expression in the IM, primordia and FMs. Signal intensity is elevated in the outermost cell layer (L1) with expression peaks at the boundary regions to emerging primordia. Expression appears to be highest in newly formed primordia and slightly attenuates during further development. (Fig. 5 A, Suppl. Fig. 7). Within the root, *MAZ* is expressed in the epidermis, particularly in root hair cells, in the vasculature, and in the RM. *MAZ* is also expressed in emerging lateral roots from early stage onwards. In mature lateral roots expression resembles the pattern in the main root (Fig 5 B, C, Suppl. Fig. 6). *MAZ* expression in the RM is characterized by elevated signal intensities in the QC and the surrounding stem cells. Expression decreases from the initials gradually towards the elongation zone. However, in developing phloem cell files fluorescence signals are not attenuated but continue upstream in the vasculature cylinder (Fig. 5 B, Suppl. Fig. 6). Marking the position of mature SEs by aniline blue staining revealed that *MAZ* expression in the differentiated vasculature is concentrated in CCs and procambial tissue, but barely in the SEs (Suppl. Fig. 1 B, C).

We detected *MAZ* signals in all *CLV1* expression domains (compare Fig. 1). Double reporter lines show co-localization of CLV1-GFP and MAZ-mCh, for example, at the border region to emerging primordia in the IM and in the distal RM (Suppl. Fig. 8). Furthermore, *MAZ* expression comprises previously reported expression domains of *BAM1* and *BAM3* (Depuydt et al., 2013; Shimizu et al., 2015). Thus, local distribution of MAZ allows its participation in different CLVf receptor pathways in Arabidopsis.

**Figure 8.**
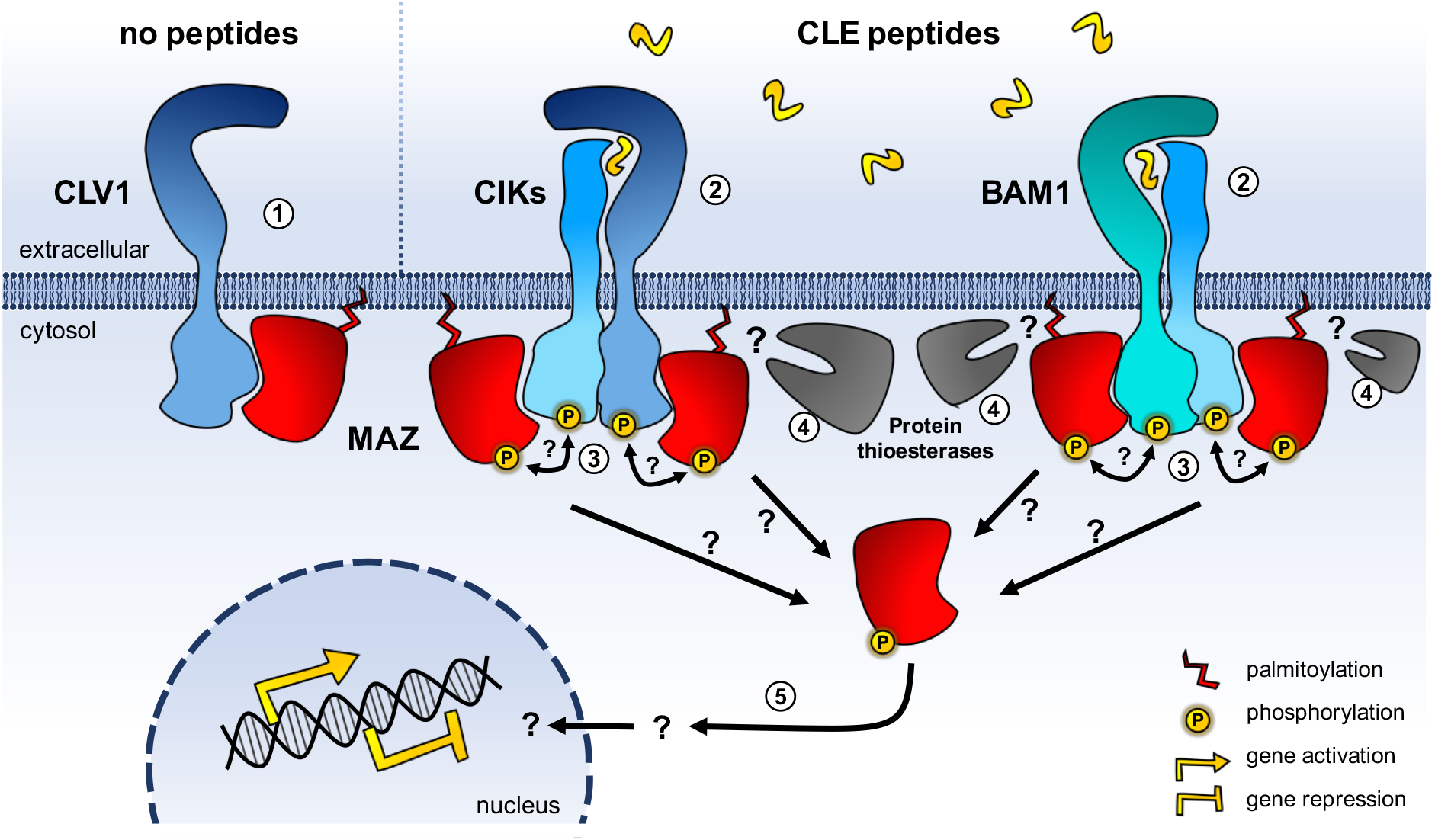
Model of MAZ as a signaling hub downstream of CLVf receptors. In the absence of CLE peptides CLVf receptors and MAZ probably build preformed complexes (**1**). The CLE peptide perception by cognate CLVf receptors is aided by CIK co-receptors (**2**). Transphosphorylation between (co-)receptors and MAZ could transmit signal downstream (**3**). Protein thioesterases could catalyze release of activated MAZ from the PM by removing palmitoylation (**4**). Activated MAZ in the cytosol could mediate signal transduction to the nucleus (**5**).

### MAZ mediates CLE peptide signaling in root development

We characterized the functional impact of MAZ within CLVf pathways by analyzing mutants harboring the T-DNA *maz* allele GABI-Kat 485F03, hereafter referred to as *maz-1*. The insertion event disrupts the KD of *MAZ* before the predicted active site and *maz-1* plants show substantial decrease of full-length *MAZ* transcripts (~30-fold in comparison to wild type samples, Suppl. Fig. 9).

Elevated organ number typically associated with *clv*-like phenotypes reflects enhanced stem cell activity in meristems, concomitant with an increase in the sizes of shoot and floral meristems. While *clv3*-9, *clv1*-20, and *crn*-10 show increased carpel numbers per silique, *maz-1* plants resemble the wild type. The double mutants clv1-20;*maz-1* and crn-10;*maz-1* resemble the single mutants *clv1-20* and *crn-10*, respectively, indicating that a loss of MAZ is insufficient to disrupt CLV3 signaling (Fig. 6 A, Suppl. Fig. 10).

To test MAZ function within CLE40-mediated stem cell differentiation in the distal RM, we quantified CSC layers in *maz-1* roots. Both, *cle40* and *clv1* mutants show increased CSC number due to reduced CLE40 signaling (Stahl et al., 2009, 2013). While increased CLE40 peptide in the growth medium reduces CSC layers in the wild type, *clv1-20* mutants are desensitized and maintain more CSC layers. *Maz-1* mutants resemble the wild type on standard growth medium but are partially resistant against external CLE40 peptide treatment that causes drastic reduction of CSC layers in Col-0 (Fig. 6 B, Suppl. Fig. 11). Notably, we found that the double mutant *clv1-20*;*maz-1* is fully sensitive to CLE40 peptide. This could point to an additional, parallel CLE40 perception pathway during CSC specification, for example via BAM1, which is activated in the absence of both CLV1 and MAZ.

CLE peptide treatment not only affects CSC fate but also root growth due to premature differentiation of the proximal RM. Disruption of CLE perception, for example, in *clv2*, *crn*, or *rpk2* mutants confers resistance against CLE-induced root shortening (Fiers et al., 2005; Shimizu et al., 2015; Müller et al., 2008). Here, we accessed root lengths of plants grown on CLV3 peptide, which is not produced in roots but can generally substitute all root-active CLE peptides in terms of root shortening, and CLE45 peptide, which is expressed in developing phloem files and perceived via BAM3 and CRN/CLV2 (Hazak et al., 2017). Our root growth assays revealed that *maz-1* single mutants are less sensitive than Col-0 seedlings to physiological concentrations of either CLV3 or CLE45 peptide. However, *maz-1* plants grown in the presence of 100 nM CLE peptides display root reductions comparable to wild type samples (Fig. 6 C). We also observed that the *maz-1* allele partially antagonizes *crn-10* mediated resistance to CLV3 peptide in *crn-10*;*maz-1* double mutants at 10 nM peptide concentration. Similar to the proposed role in CSC fate control, this could again indicate a dual function of MAZ to mediate CLE responses and simultaneously repressing parallel acting CLE perception pathways.

Together, our data suggest that MAZ is involved in several CLE signaling pathways, involving CLV1 and CLV2/CRN.

We also accessed morphology and physiological constraints of *maz-1* plants regarding previously described phenotypes of other *pti1-like* mutants, i.e. impaired root hair development of mri mutants (Boisson-Dernier et al., 2015), and hypersensitivity to drought stress like *cark1* and *cark6* mutants (Zhang et al., 2018; Wang et al., 2019). However, in contrast to *mri-2* and *fer-4* mutants, *maz-1* seedlings display mean root hair length at wild type-level (Suppl. Fig. 12). Furthermore, within two different experimental setups, neither *maz-1* mutants nor *MAZ* overexpression lines show altered drought stress responses, which have been associated with disturbed ABA signaling of mutants or overexpressors of *CARK1* and *CARK6*, respectively (Suppl. Fig. 13, 14).

### A novel role for CLV1 with MAZ in stomatal spacing

Regular patterning of stomata at the leaf epidermis is the result of finely controlled series of cell divisions of the stomata cell lineage. In this process meristemoid mother cells (MMCs) divide asymmetrically to form a meristemoid and a larger sister cell. While meristemoids are precursors of guard mother cells (GMCs) that differentiate into the two GCs of a stoma, the sister cell may become a pavement cell or undergoes a spacing division before generating a new meristemoid (reviewed in (Bergmann and Sack, 2007)). Mutants lacking this regular spacing between two stomata, for example, due to disruption of the receptor complex that senses positional cues in the leaf epidermis, include, among others, *er;erecta-like1(erl1);erl2* triple mutants of the *ERf* receptor genes and mutants of the LRR-RLP encoding *TOO MANY MOUTHS (TMM)* gene. These mutants accumulate two or more stomata directly adjacent to each other, i.e. display stomatal clustering (reviewed in (Zoulias et al., 2018)).

We observed that *clv1* mutants are also associated with defects in regular patterning in the leaf epidermis (Suppl. Fig. 15). Initial data revealed an increased number of stomata in true leaves, but not in cotyledons, of clv1-20 plants in comparison to wild type Col-0, uncovering a novel role for CLV1 in stomatal patterning. Furthermore, CLE peptide treatment antagonized this effect in *clv1-20* mutants (Suppl. Fig. 15). We extended these analyses to include *maz-1* and the double mutant *clv1-20*;*maz-1*. Quantification of stomatal clusters in seedlings 14 DAG grown on ½ MS agar plates, not only confirmed defects in patterning of *clv1-20* true leaves, but also showed that the double mutant *clv1-20*;*maz-1* enhances this phenotype (Fig. 7). In comparison to the wild type control (Col-0) the mean stomata cluster rate of *clv1-20* is elevated (~ 2 % stomata in cluster). However, in *clv1-20*;*maz-1* cluster rate is significantly increased to ~ 10 %, while the single *maz-1* mutant displays no clusters. Stomata density was not significantly different between genotypes. This points towards a specific function of CLV1 and *MAZ* in spacing divisions, but not stomata specification. Since both, *MAZ* and *CLV1*, are expressed in cells of the stomatal lineage (Fig. 1 E, 5 D), we propose that the CLV1/MAZ signaling module serves to establish stomata spacing.

## Discussion

CLVf RLKs participate in various signaling systems, thereby controlling meristem homeostasis but also processes like vascular formation or plant immunity. Here, we add stomata development to the scope of CLV1 functions. Considering the diversity of CLVf pathways, mechanisms to mediate specific downstream responses are required. In their role as direct interactors of CLVf receptors, MAZ and several of its homologs are prime candidates to participate in downstream signaling after CLE perception. Furthermore, the expression pattern of *MAZ* overlaps with the places of action of the CLVf receptors in Arabidopsis. Therefore, MAZ might integrate and cross-regulate signaling cues from different CLVf receptors.

### Functional redundancy of Pti1-like homologs could mask mutant phenotypes

We found that *maz-1* plants are resistant to exogenous CLE treatment in the context of CSC specification (Fig. 6 B). However, in comparison to *clv1-20* mutants, the resistance of *maz-1* is less pronounced. Also, the CLE peptide resistance of *maz-1* seedlings regarding proximal RM development is less penetrating than in mutants of the involved receptors. At higher CLE peptide concentrations root shortening of *maz-1* seedlings resembles the level of wild type-samples (Fig. 6 C). Furthermore, the *maz-1* allele confers no *clv1*-like increased carpel number (Fig. 6 A). These observations suggest that other Pti1-like homologs compensate MAZ function in *maz-1* mutants. In line, several Pti1-like proteins interact with CLV1, reflecting a conserved binding capacity of Pti1-like homologs to the CLVf receptors (Fig. 3 C).

Functional redundancy is a common feature within the Pti1-like family. For example, Pti1-1, Pti1-2, and Pti1-3 (=MAZ), respectively, interact with the stress-related kinase OXI1 (shown by yeast-two-hybrid and Co-IP). However, only the interaction of OXI1 with Pti1-2 was validated by stimulus triggered phosphorylation *in vivo* (Anthony et al., 2006). This indicates a high general potential of functional redundancy but at the same time demonstrates the presence of mechanisms that add specificity to the respective Pti1-like homologs within their physiological context.

### The N-terminus of Pti1-like proteins could determine functional specification

The 11 Pti1-like homologs in Arabidopsis are especially conserved within the KDs, while their more variable N-termini allow specification (Appendix). As we demonstrated in here, the MAZ N-terminus determines its subcellular localization via palmitoylation (Fig. 4). In contrast to other mechanisms that mediate PM-localization, the attachment of a palmityl group via S-acylation is reversible. As such, palmitoylation can dynamically regulate the subcellular localization of proteins (reviewed in (Hurst and Hemsley, 2015)). All Pti1-like family members in Arabidopsis, except Pti1-6, harbor conserved cysteines, which are predicted palmitoylation sites (Suppl. Tab. 3). In general, S-acylation is often coincidently found with protein myristoylation. However, within the Arabidopsis Pti1-like family only CARK1 is additionally equipped with a myristoylation site, suggesting it is obligatorily PM-localized. The subcellular localization of the other nine Pti1 homologs in Arabidopsis might be dynamically controlled by counteracting protein S-acyl transferases (PATs) and acyl protein thioesterases. Although the underlying mechanisms in plants are not well understood (reviewed in (Li and Qi, 2017)), palmitoylation allows stimulus-triggered subcellular re-localization and display a powerful tool for signal transduction. S-acylation of Pti1-like proteins is evolutionary conserved among various species, including monocots, as shown e.g. for ZmPti1a from maize and OsPti1a from rice (Herrmann et al., 2006; Matsui et al., 2014).

Notably, the MAZ N-terminus is longer than in most of its Arabidopsis homologs, potentially offering additional target sites for specific control, such as differential phosphorylation or protein turnover via ubiquitination. It comprises two Phospho-Serines (detected in the PhosphAt Database (Heazlewood et al., 2008)) and according to different prediction tools 3 – 7 lysine residues as potential targets for ubiquitination.

### CLV1/MAZ contributes to spacing divisions in stomata development

Feedback-regulated signaling systems are crucial to integrate positional information from neighboring cells or tissues into developmental processes and to ensure dynamic but controlled growth. Such feedback loops determine, for instance, the stem cell domain in the SAM, IMs, and FMs (Fletcher et al., 1999; Brand et al., 2000), and cell fate in the leaf epidermis during stomatal patterning (reviewed in (Tameshige et al., 2017)). While the CLV pathway is the major regulative element in the shoot meristems, the ERf receptors are essential for cell fate decisions during leaf epidermis differentiation. However, the ERf also impacts stem cell homeostasis in the shoot meristems (Uchida et al., 2013; Mandel et al., 2014; Zhang et al., 2020). In turn, CLV-related pathways are involved in stomatal development, e.g. by perceiving CLE9/10 peptides (Qian et al., 2018). In here we provide first evidences that *clv1-20* mutants are defective in regular patterning of GCs and that *maz-1* enhances this phenotype as a second site mutation (Fig. 7).

In the epidermis of cotyledons and true leaves the *CLV1* promotor is predominantly active in meristemoids and stomatal lineage ground cells, while *CLV1* expression is barely detectable in differentiated epidermis cells, neither in pavement cells nor in GCs (Fig. 1). This hints toward a specific function of the CLV1 receptor within the precise signaling events of leaf epidermis differentiation. The notion that *clv1* mutants, in contrast to, for example, *tmm* mutants (Geisler et al., 2000), show rather mild stomatal patterning defects suggests the presence of redundantly acting genes. Furthermore, stomata density in *clv1* mutants is comparable to the wild type (Fig. 7). Thus, CLV1 does not determine MMC fate, but may act later in the stomatal lineage to establish the one-cell-spacing rule between two stomata. This stomatal spacing is mainly regulated by EPIDERMAL PATTERNING FACTOR 1 (EPF1) and its primary receptor ERL1, which form a negative feedback loop with the transcription factor MUTE (Hara et al., 2007; Lee et al., 2012; Qi et al., 2017). CLV1-based signaling could interfere with this feedback regulation or act in parallel to finetune spacing divisions.

Our observation that the double mutant *clv1-20*;*maz-1* displays increased stomata cluster rate in comparison to *clv1-20* single mutants (Fig. 7) suggests that MAZ is not only involved in stomatal pattering as a potential downstream target of CLV1, but also affects spacing divisions through CLV1-independent signaling. Therefore, *MAZ* is possibly a shared downstream target of CLV1 and other RLKs, e.g. ERL1. Since *maz-1* single mutants are wild type-like regarding stomata clusters, redundancy mediated by other Pti1-like homologs can be assumed. However, in combination with the *clv1-20* mutation this redundancy is partly overcome. This might be explained by the quantitative character of CLV1/MAZ signaling during stomatal development comprising additional, yet unknown components.

## Conclusions

We here report on the identification of MAZ and other RLCKs of the Pti1-like family as intermediators of CLVf receptor responses. Within the versatile functional contexts of CLVf pathways, MAZ and redundantly acting homologs could contribute to differential signaling outputs. After CLE perception by CLVf receptors, signal transduction may involve transphosphorylation between the CLVf RLKs and Pti1-like RLCKs and subsequent detachment of MAZ or its homologs from the membrane to reach intracellular targets (compare Fig. 8). Deciphering how MAZ and the CLVf are functionally connected will be critical for further characterization of the Pti1-like family as downstream elements of CLVf receptor pathways.

## Material and Methods

Detailed information on chemicals used for the described experiments are available in Suppl. Tab. 4.

### Plant material and growth conditions

All *Arabidopsis thaliana* (L.) Heynh. plants in this study are ecotype Columbia-0 (Col-0). Origin and details on utilized mutants harboring the alleles *maz-1*, *clv1-20*, *crn-10*, *clv3-9*, *cark1-2*, *cark6-1*, *mri-2*, and *fer-4*, respectively, can be found in Suppl. Tab. 5. The presence of the respective alleles was controlled following genotyping strategies as listed in Suppl. Tab. 6. Before sowing, *A. thaliana* seeds were sterilized in ethanol solution (10 min in 80 % v/v ethanol, 1.3 % w/v sodium hypochlorite, 0.02 % w/v SDS), or in a chloric gas atmosphere (1 h in a desiccator after mixing 50 ml of 13 % w/v sodium hypochlorite with 1 ml 37 % HCl,). If not indicated differently, plants were cultivated in phytochambers under long day (LD) conditions (16 h light / 8 h dark) at 21 °C on soil. Alternatively, seedlings were cultivated for up to 14 days on ½ MS agar plates (1 % w/v sucrose, 0.22 % w/v MS salts + B5 vitamins, 0.05 % w/v MES, 12 g/l plant agar, adjusted to pH 5.7 with KOH) in phytocabinets (continuous light, 60 % humidity, and 21 °C).

*N. benthamiana* plants were grown 4 – 5 weeks in the greenhouse and subsequently used for transient leaf epidermis cell transformation. After infiltration, *N. benthamiana* plants were kept under high humidity in continuous light.

### Cloning

Information on the entry plasmids, subsequently used for the assembly of plant expression vectors can be found in Suppl. Tab. 7. New entry plasmids generated in this study contain DNA sequences that were PCR-amplified (with Phusion High-Fidelity PCR polymerase) from genomic DNA prepared of Col-0 rosette leaves. PCR fragments were introduced via customized oligonucleotide overhangs to either pENTR^®^ (Gateway^®^ system, (Katzen, 2007)) or pGGA000 and pGGC000 (GreenGate system, (Lampropoulos et al., 2013)). Coding sequences of genes of interest (GOIs) were amplified without the STOP codon to allow C-terminal fusions with fluorophores. Single positions within gene sequences were modified by site-directed mutagenesis applying the QuikChange II kit according to manufacturer’s protocol (Agilent Technologies).

Destination plasmids utilized for transient expression in *N. benthamiana* and for generation of stable *A. thaliana* lines are listed in Suppl. Tab. 8 and 9, respectively. Inducible vectors were created by a Gateway^®^ LR reaction (according to manufacturer’s instructions, Thermo Fisher Scientific), combining an entry vector harboring the GOI with either pABindGFP or pABindmCherry (Bleckmann et al., 2010). Transgenes for constitutive *N. benthamiana* and stable Arabidopsis expression were constructed with the GreenGate system, assembling desired DNA sequences and the backbone pGGZ001 following a golden gate principle (compare Suppl. Tab. 8 and 9).

To amplify plasmid DNA, competent Escherichia coli DH5α cells were heat-shock transformed and cultivated on selective LB medium (1 % w/v tryptone, 0.5 % w/v yeast extract, 0.5 % w/v NaCl). After plasmid DNA purification via commercial kits, the plasmids were validated by restriction digest and Sanger sequencing.

### Transient gene expression in *N. benthamiana*

Agrobacterium-mediated transformation of leaf epidermis cells of *N. benthamiana* was applied to monitor transient expression of transgenes. The *Agrobacterium tumefaciens* strain GV3101 pMP90 (rifampicin and gentamycin resistant) was used. Bacteria were transformed with the required plasmid vectors via the heat-shock method (aliquots of competent cells mixed with 1 μl 100 nM plasmid DNA were incubated for 5 min in liquid N_2_ followed by 5 min at 37 °C and subsequent regeneration). All Gateway^®^-based destination plasmids were introduced to GV3101 pMP90 additionally equipped with an expression cassette for the p19 suppressor of gene silencing from the tomato bushy stunt virus to enhance efficiency of transgene expression (Voinnet et al., 2003). Plasmids constructed via the GreenGate system were introduced to *A. tumefaciens* GV3101 pMP90 pSoup. The helper plasmid pSoup confers resistance against tetracycline and contains the RepA gene, which encodes a trans-activating replicase for the pSA origin of replication that is mandatory for propagation of GreenGate-based destination plasmids in *A. tumefaciens* (Hellens et al., 2000).

For infiltration of *N. benthamiana* leaves, agrobacteria containing the respective expression cassettes were cultivated overnight with shaking at 28 °C in 5 ml dYT (double Yeast Tryptone, 1.6 % w/v tryptone, 1 % w/v yeast extract, 0.5 % w/v NaCl) with appropriate antibiotics (Gateway^®^ plasmids in GV3101 pMP90 p19 with 50 μg mL^−1^ rifampicin, 50 μg mL^−1^ gentamycin, 50 μg kanamycin, and 100 μg mL^−1^ spectinomycin; GreenGate plasmids in GV3101 pMP90 pSoup with 50 μg mL^−1^ rifampicin, 50 μg mL^−1^ gentamycin, 2.5 μg tetracycline, and 100 μg mL^−1^ spectinomycin). Cell cultures were adjusted to an optical density (OD_600nm_) of 0.3 and centrifuged (10 min, 4000 x g, 4 °C). The pellet was resuspended in infiltration medium (5 % w/v sucrose, 150 μM acetosyringone, 0.01 % v/v Silwet) and incubated at 4 °C for 2 – 3 h. For co-expression of two or more transgenes, the corresponding transformed *A. tumefaciens* strains were mixed equally (final OD_600nm_ = 0.3 per strain). Subsequently, the bacterial resuspensions were infiltrated with a syringe into the abaxial site of the *N. benthamiana* leaves.

Plants transformed with constructs under the control of an estradiol-inducible promotor system (Zuo et al., 2000) were sprayed 2 – 4 days after infiltration and 6 – 16 h prior to sample preparation for imaging or CoIP with an estradiol solution (10 μM ß-estradiol, 0.1 % v/v Tween-20). Constitutive expressing transgenes (under the control of the *UBQ10* promotor) were used for analyses 3 days after infiltration.

### Stable transformation of *A. thaliana*

To generate stable expression lines, parental *A. thaliana* plants were transformed 4 – 6 weeks after germination via the floral dip method using *A. tumefaciens* GV3101 pMP90 pSoup previously transformed with desired binary vectors (Suppl. Tab. 9, (Clough and Bent, 1998)). Agrobacteria were cultivated overnight (28°C, shaking) in selective dYT medium (supplied with 50 μg mL^−1^ rifampicin, 50 μg mL^−1^ gentamycin, 2.5 μg tetracycline, and 100 μg mL^−1^ spectinomycin). 50 ml main cultures of the transgene-harboring Agrobacterium strains were inoculated with 1 ml of an overnight preculture and centrifuged the next day (10 min, 4000 x g, 4 °C). The pellet was resuspended in transformation medium (5 % w/v sucrose, 10 nM MgCl_2_, 0.01 % Silwet). Plants were prepared by removing already developed siliques and were then subjected to floral dip. Whole shoots with special attention to the flowers were immersed into the *A. tumefaciens* resuspension for 30 sec up to 5 min to guarantee the entry of bacteria into the floral tissue, more precisely the female gametophyte. Afterwards, the plants were kept under high humidity overnight at room temperature. Subsequently, the plants were cultivated in standard phytochamber conditions until harvest. The t1 seeds were screened for positive transformants by selection either for resistance against DL-phosphinothricin (PPT, alternatively used in form of the herbicide BASTA^®^), or against hygromycin. Seeds of the transformed t0 plants were screened for positive transformants by selection either for resistance against DL-phosphinothricin (PPT, alternatively used in form of the herbicide BASTA^®^), or against hygromycin. T1 plants harboring transgenes with the BASTA^®^ resistance cassette were selected by spraying seedlings on soil (10 DAG) with a 120 mg/ml solution of BASTA^®^, or by sowing the t1 seeds on ½ MS agar plates supplied with 10 μg/ml PPT. Transgenic plants with hygromycin resistance were screened on ½ MS agar plates supplied with 15 μg/ml hygromycin. Positive plants were further amplified for t2 selection and identification of homozygous lines. All stable *A. thaliana* lines used and generated in this work are listed in Suppl. Tab. 10.

### Confocal microscopy and tissue staining

*In vivo* fluorescence microscopy was performed at the CLSM systems Zeiss LSM 780 and Zeiss LSM 880, respectively, employing C-Apochromat 40x/1.20 water objectives. For whole leaf imaging and quantifying stomata (Fig. 7, Suppl. Fig. 16) a Plan-Apochromat 10x/0.45 M27 air objective was used. Samples containing GFP derivates were excited with an argon laser at 488 nm and emission was detected at 490 – 530 nm by a 32-channel GaAsP detector or the Airyscan detector system with a BP 495-550 / BP 570-620 filter set. The argon laser was also used for excitation of mVenus at 514 nm, followed by measuring the emission at 520 – 550 nm with a GaAsP detector. Diode-pumped solid state (DPSS) lasers were employed to excite mCh at 561 nm. Emission was detected by photomultiplier tubes (PMTs) in the range of 570 – 650 nm. Propidium iodide (PI) was utilized to stain cell walls in roots (25 μM), shoot meristems (5 mM), and leaf epidermis (50 μM). To quantify CSC layers, roots were subjected to mPS-PI staining according to (Truernit et al., 2008), thereby marking cell walls and starch granules in differentiated columella cells. PI was excited at 561 nm by DPSS lasers and detected by PMTs at 590 – 650 nm. Alternatively, cell walls were counterstained with DAPI, which was excited at 405 nm with a laser diode and emission was recorded at 410 – 480 nm by PMTs. To visualize sieve plates of SEs, roots of Arabidopsis seedlings were incubated for 5 min in 0.01 % w/v aniline blue, subsequently rinsed in a washing solution (10 mM KCl, 10 mM CaCl_2_, 5 mM NaCl), and imaged (excitation with a 405 nm diode, emission detected at 470 – 530 nm with PMTs).

### FRET-FLIM interaction analysis

FRET-FLIM experiments were conducted at a Zeiss LSM 780 (C-Apochromat 40x/1.20 water objective) equipped with a single-photon counting device (PicoQuant Hydra Harp 400) and a linear polarized diode laser (LDH-D-C-485). GFP or mNeonGreen donor fluorophores were excited at 485 nm with a pulsed laser at a frequency of 32 MHz. Excitation power was adjusted to 1 μW. Emission was detected in perpendicular and parallel polarization by Tau-SPADs (PicoQuant) with a band-pass filter (520/35 AHF). Images were acquired with a frame size of 256 × 256, zoom 8, and a pixel dwell time of 12.6 μs. For each measurement 60 frames were taken and the intensity-weighted mean lifetimes τ [ns] were calculated using PicoQuant SymPhoTime64 software applying a biexponential fit. The displayed data were obtained from at least 3 independent experiments.

### Protein extraction and CoIP

Plant material was collected and immediately frozen in liquid N_2_. All following steps were performed at 4 °C. To identify novel CLV1 interactors via CoIP, whole Arabidopsis seedlings were used (grown in liquid ½ MS, continuous light, gently shaking). Medium was removed 7 DAG and plants were grinded in liquid N_2_ with mortar and pestle to obtain fine powder. Per sample 500 mg material was mixed with 750 μl extraction buffer (EB: 50 mM Tris/HCl pH 7.5, 150 mM NaCl, 1 mM EDTA, 10 % v/v glycerol, 0.1 % v/v Nonident P40 substitute, 5 μM dithiothreitol (DTT), 1 tablet of cOmplete™ proteinase inhibitor cocktail dissolved in 50 ml EB). For interaction assays, 6 leaf discs (6 mm diameter, ~ 300 mg fresh weight) from transiently transformed *N. benthamiana* leaves were grinded in a tube with two glass beads (3 mm) with a TissueLyser II (Qiagen) and directly after supplied with 600 μl EB. Samples dissolved in EB were incubated on a rotator for 1.5 h and subsequently centrifuged (20 min, 17 x g). The supernatant was collected for immunodetection via sodium dodecyl sulfate polyacrylamide gel electrophoresis (SDS-PAGE, standard protocol of NEXT-GEL^®^ with 10 % acrylamide, samples were mixed with loading buffer (5 x: 200 mM Tris/HCl pH 6.8, 8 % w/v SDS, 40 % v/v glycerol, 0.05 % w/v bromophenol blue, 50 mM DTT), and boiled 5 min at 95 °C prior to loading) and Western Blot (WB, wet electroblotting system, 80 min / 100 V) analysis or used as input material for CoIP.

For CoIP experiments 500 μl protein extract was mixed with 25 μl calibrated anti-GFP magnetic beads (ChromoTek GFP-Trap^®^) and incubated on a rotator for 2 h. After, supernatant was discarded and beads were washed 4 times with washing buffer (10 mM Tris/HCl pH 7.5, 150 mM NaCl, 0.5 mM EDTA). Loaded beads were subjected to mass spectrometry or proteins were eluted from beads by mixing with double concentrated loading buffer and boiling (95 °C / 5 min) for analysis via SDS-PAGE, WB, and alkaline phosphatase (AP)-based immunostaining. To detect GFP-fusion proteins, primary GFP antibodies (IgG, rat monoclonal, diluted 1:1500 in blocking solution, incubation overnight at 4°C) and secondary anti-rat-AP antibodies (1:5000, 2.5 h, room temperature) were applied. To detect RFP derivates, primary RFP antibodies (IgG, mouse monoclonal, 1:1500, overnight, 4 °C) and secondary goat anti-mouse IgG AP antibodies (1:5000, 2.5 h, room temperature) were used.

### Mass spectrometric analysis

Mass spectrometry was performed at the Molecular Proteomics Laboratory (MPL, BMFZ, HHU Düsseldorf). After immunoprecipitation of samples with anti-GFP beads (free GFP n=5, CLV1-2xGFP n=5, CLV1-2xGFP + 5μM CLV3 peptide n=4), precipitated proteins were eluted with 25 μl sample buffer (7.5% v/v glycerine, 3% w/v SDS and 37.5 mM 2-Amino-2- (hydroxymethyl)-1,3-propanediol in water, pH 7) for 10 minutes at 90°C. Samples were further processed by in-gel digestion with trypsin including reduction with dithiothreitol and alkylation with iodoacetamide as described (Grube et al., 2018). Resulting peptides were resuspended in 0.1% trifluoroacetic acid and about half of the sample from each immunoprecipitation analyzed by liquid chromatography coupled mass spectrometry as previously described (Ingold et al., 2018). Briefly, peptides were separated on C18 material on an Ultimate 3000 rapid separation liquid chromatography system (Thermo Fisher Scientific) using a one-hour gradient and injected into a QExactive plus mass spectrometer (Thermo Fisher Scientific) via a nano electrospray source interface. The mass spectrometer was operated in data dependent, positive mode. First, survey scans were recorded (resolution 70000, scan range 200 – 2000 m/z) and subsequently, up to twenty > 1 charged precursors ions were selected by the build-in quadrupole, fragmented by higher-energy collisional dissociation and MS/MS spectra recorded at a resolution of 17500.

Data analysis was carried out with MaxQuant (version 1.6.0.16, Max Planck Institute for Biochemistry, Planegg, Germany) with standard parameters if not stated otherwise and using the *A. thaliana* reference proteome sequences (UP000006548, downloaded on 2nd February 2017 from UniProt) supplemented with two entries for CLV1-2xGFP and free GFP. The “match between runs” function and label-free quantification was enabled. Peptides and proteins were accepted for identification with a false discovery rate of 1%. Only proteins identified with at least two different peptides were considered as identified. Precursor intensity-based quantification data was further analyzed using the Perseus framework (version 1.6.0.7, Max Planck Institute for Biochemistry, Planegg, Germany).

### Reverse transcriptase quantitative real-time PCR (RT-qPCR)

To evaluate the amount of residual *MAZ* transcripts in *maz-1* plants, RNA from seedlings 7 DAG was extracted with the Qiagen RNeasy Plant mini kit and 2 μg RNA per reaction was used for first strand cDNA synthesis via SuperScript™ III reverse transcriptase (according manufacture’s protocol with oligo(dT)_18_). The qPCR reactions were performed with SsoAdvanced™ Universal SYBR^®^ Green Supermix in a Stratagene Mx3005P qPCR System (Agilent Technologies) with an optimized dilution of cDNA (accessed via dilution series for each oligo pair). Applied oligonucleotides are listed in Suppl. Tab. 11. Data were normalized to the housekeeping genes AT2G28390, AT4G34270, and At4g26410. Technical triplicates of 3 biological replicates were considered for each condition.

### Evaluation of mutant morphology

Carpel number per silique of different *A. thaliana* mutants grown on soil under LD and continuous light conditions, respectively, were quantified from 10 – 15 plants per genotype and 15 – 30 siliques per plant. Root length of seedlings 10 DAG was accessed after cultivation of the examined genotypes on ½ MS plates supplied with synthetic CLV3 peptide in the indicated concentrations. Measurements were done with ImageJ after scanning the plates. For each condition at least 67 (and up to 113) single roots were measured and normalized to the mean of the Col-0 samples of the same peptide concentration.

Root hair length and density of seedlings 5 DAG were quantified after cultivation on ½ MS and image acquisition with a Nikon SMZ25 stereomicroscope. Per genotype 20 – 25 roots were analyzed, in total with 3542 (Col-0), 975 (*fer-4*), 2433 (*mri-2*), and 4360 (*maz-1*) single measurements. For each seedling, the length of all root hairs in focus on one side of the root were determined.

Stomata cluster in cotyledons and leaves of seedlings 14 DAG grown on ½ MS were analyzed after PI staining and imaging by counting total number of stomata and abundance of directly adjacent stomata. For each genotype 5 – 10 cotyledons and 6 – 10 true leaves were examined by counting all stomata in 4 distinct regions (400 x 400 μm) of each sample.

### Evaluation of mutant physiologic responses

Drought stress experiments to access potential involvement of the *maz-1* allele in ABA responses were performed with plants of different genetic background grown on soil under standard conditions. Two weeks after germination seedlings were subjected to water deficiency for either 10 days or 20 days and were subsequently re-watered to monitor recovery. Both, the 10 days and 20 days drought period approaches, included 3 pots of plants for each genotype, which were randomly distributed on the tray.

Water-loss assays were conducted to estimate the degree of evaporation via open stomata after dissection of 6 rosette leaves per sample (1 sample = 1 individual plant). For each genotype 8 samples (per repeat) were weighted at indicated time points. The reduction of weight was taken as an approximation for water-loss. The assay was performed in three independent repeats.

### Software

Statistical analyses and data plotting were realized with GraphPad Prism v 8. For visualization and quantification of microscopic data ZEN (Zeiss, Black Version) and ImageJ v 1.51 (Schneider et al., 2012) was employed. The following prediction tools were used: CSS-Palm 4.0 (Ren et al., 2008), PredGPI (Pierleoni et al., 2008), Myristoylator by ExPASy (Bologna et al., 2004), UbPred (Radivojac et al., 2010), UbiSite (Huang et al., 2016), BDM-PUB v 1.0. Cloning was organized via Vector NTI^®^ software. Protein alignments were done with ClustalΩ (Sievers et al., 2011), phylogenetic analyses with MEGA X (Kumar et al., 2018), and tree visualization with iTOL v 4 (Letunic and Bork, 2019).

## Acknowledgements

P.B. was supported by the IMPRS *Understanding complex plant traits using computational and evolutionary approaches*, work in R.S. lab is supported by the DFG through CRC1208 and CEPLAS (EXC 2048). We thank Cornelia Gieseler, Silke Winters, and Carin Theres for technical support, Aurélien Boisson-Dernier and Zachary Nimchuk for sharing Arabidopsis seeds, Grégoire Denay, Rebecca Burkart, Marc Somssich, Frederic Boyer, and Pauline Anne for sharing plasmids, Dolf Weijers and Mark Roosjen for the “sticky list” of typical GFP control pull-downs, Vicky Howe for critical reading of the manuscript, the Center for Advanced imaging (CAi) at HHU for microscopy support, and Gereon Poschmann from the Molecular Proteomics Laboratory (MPL) at HHU for mass spec analyses.

## Authors contributions

R.S. and P.B. conceived and planed the project. P.B. conducted experiments and data analyses, besides the following: J.S. performed shoot meristem imaging and image processing, S.B. did FLIM interaction studies of CLV1-mNeonG vs. Pti1-like homologs (Fig. 3 C), K.P. acquired initial stomatal cluster rate data (Suppl. Fig. 15). P.B. and R.S. wrote the manuscript.

## Supplementary material

**Supplemental Figure 1** Expression domains of *CLV1* and *MAZ* in the vasculature of Arabidopsis roots.

**Supplemental Figure 2** Two independent CoIP repetitions of CLV1-GFP with MAZ-mCherry.

**Supplemental Figure 3** Subcellular localization of MAZ in *N. benthamiana* leaf epidermis cells.

**Supplemental Figure 4** Predicted palmitoylation sites at the N-terminus of MAZ are mandatory for PM-localization.

**Supplemental Figure 5** Expression pattern of the transcriptional reporter *MAZ:mVenus-NLS*//Col-0 in the root (3 DAG).

**Supplemental Figure 6** MAZ expression by MAZ promotor in different translational reporters.

**Supplemental Figure 7** Expression of *MAZ:MAZ-eGFP* in the SAM of 5 weeks old *A. thaliana*.

**Supplemental Figure 8** Co-localization of CLV1-eGFP and MAZ-mCherry in shoot and root.

**Supplemental Figure 9** Characterization of the *maz-1* allele GABI-Kat 485F03.

**Supplemental Figure 10** Carpel number of *maz-1* mutants is wild typic in continuous light (CL).

**Supplemental Figure 11** *Maz-1* mutants are partially resistant to CLE40 peptide treatment in terms of columella stem cell (CSC) layer specification.

**Supplemental Figure 12** The *maz-1* mutant does not display defects in root hair development.

**Supplemental Figure 13** Water loss assay of Arabidopsis leaves to monitor potential differential stomata closure as an indication for a role of Pti1-homologs in ABA signaling.

**Supplemental Figure 14** Drought stress assay revealed no clear differences between the analyzed genotypes to cope with water deficiency.

**Supplemental Figure 15** Impact of the *clv1-20* allele and CLE40 peptide treatment on stomata development and clustering in Arabidopsis seedlings.

**Supplemental Figure 16** Cotyledons of *clv1-20*, *maz-1*, and the double mutant *clv1-20;maz-1* display no significant changes of stomata density and cluster rate.

**Supplemental Table 1** Via mass spectroscopy identified proteins in CoIP fraction against CLV1-2xGFP, which are not found in GFP-only control samples.

**Supplemental Table 2** Mean FRET efficiencies of tested donor-acceptor combinations at the PM of transiently transformed *N. benthamiana* epidermis cells.

**Supplemental Table 3** Pti1-like family in *Arabidopsis thaliana* and predicted PM-localization mechanisms.

**Supplemental Table 4** Chemicals.

**Supplemental Table 5** Analyzed *A. thaliana* mutants (Col-0 background).

**Supplemental Table 6** Genotyping strategies to verify listed mutant alleles.

**Supplemental Table 7** Entry plasmids.

**Supplemental Table 8** Plasmids used for transient gene expression in *N. benthamiana*.

**Supplemental Table 9** Plasmids used for stabile transformation of *A. thaliana*.

**Supplemental Table 9** Transgenic *A. thaliana* lines applied and generated in this study.

**Supplemental Table 11** Oligonucleotides for RT-qPCR.

**Appendix** Sequence alignment with ClustalΩ of Arabidopsis Pti1-like family and SlPti1.

## Notes

### Competing Interest Statement

The authors have declared no competing interest.

