## Supplemental Figures 1 - 16 for "The receptor-like cytoplasmic kinase MAZZA and CLAVATA-family receptors interact *in vivo*, together mediating developmental processes in *Arabidopsis thaliana*"

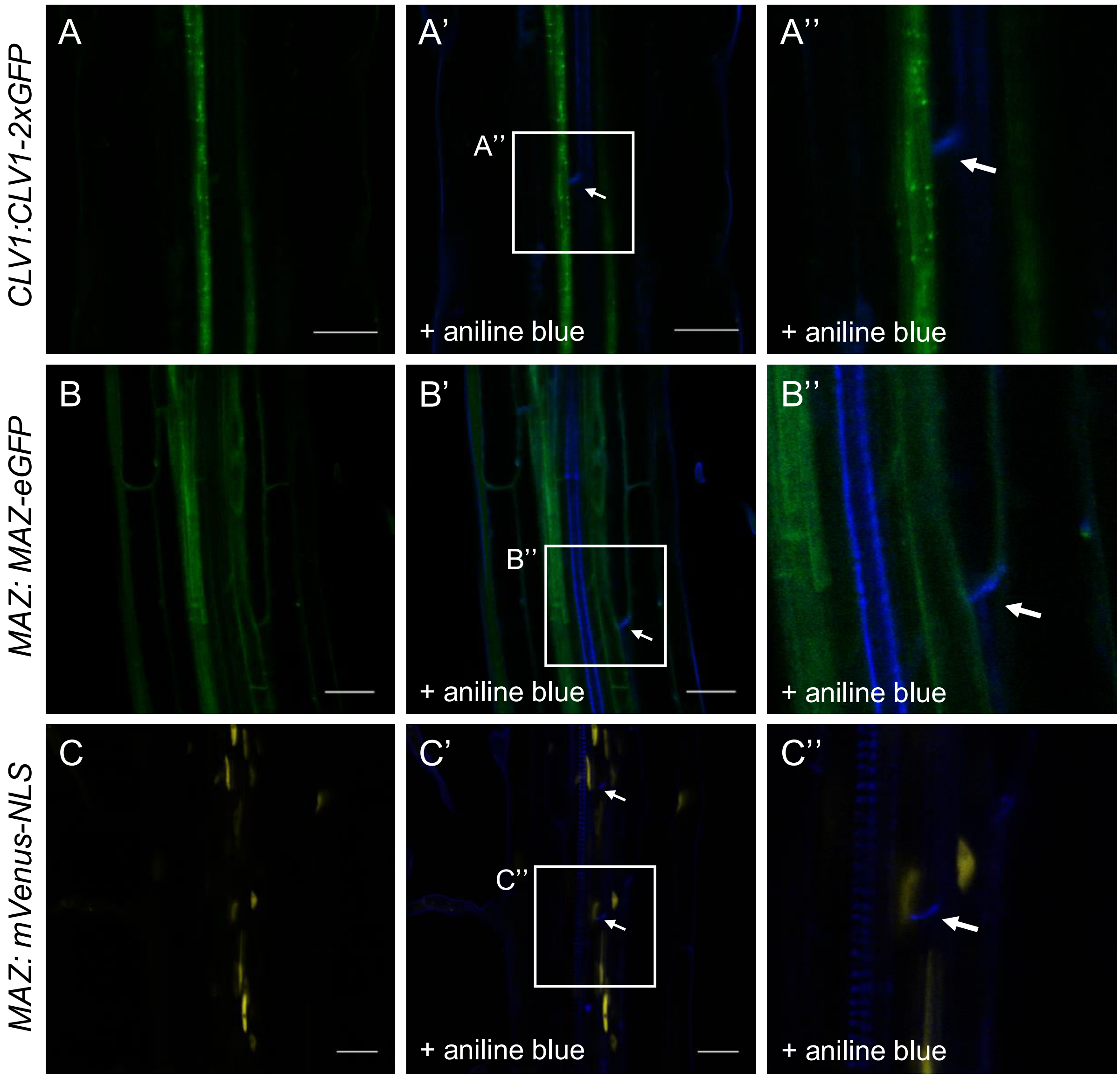


**Supplemental Figure 1** Expression domains of *CLV1* and *MAZ* in the vasculature of Arabidopsis roots.

Different reporter lines 3 DAG grown on ½ MS were counterstained with aniline blue to visualize callose at the sieve plates (white arrows) and thereby marking the position of sieve elements (SE) within the vasculature cylinder. **A – A''** The *CLV1-2xGFP* fusion under the control of the endogenous *CLV1* promotor is expressed in companion cells (CC), directly adjacent to the SEs, which show no fluorescence signal. **B – B''** The *MAZ:MAZ-eGFP* reporter also mediates strong expression in the CCs. However, the expression domain is extended to other cell files in the vasculature, including the procambial tissue. In contrast, signals from the SEs are barely detectable. **C – C''** Expression pattern of the transcriptional *MAZ:mVenus-NLS* reporter in the vasculature cylinder are in line with the translational reporter in B – B''. Scale bars: 25 µm.

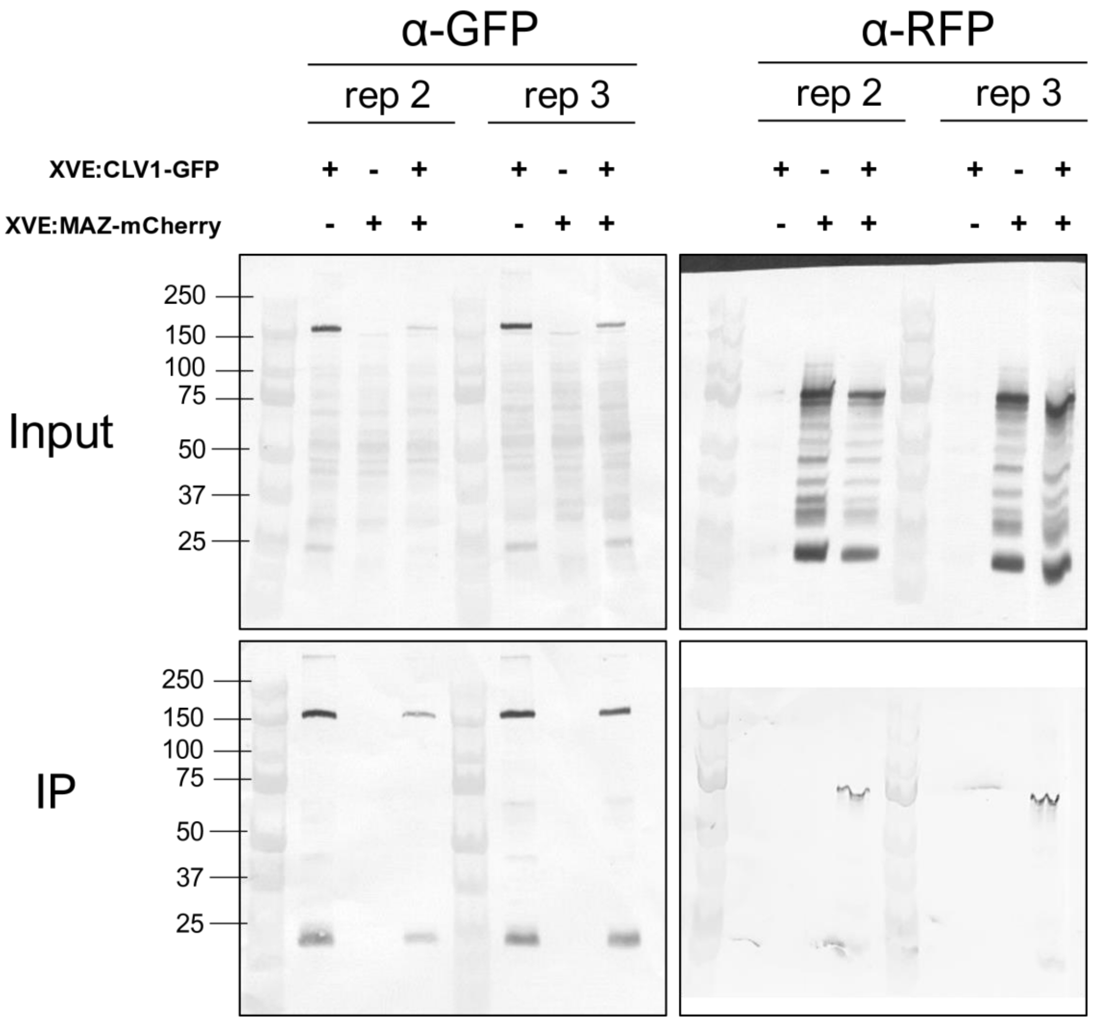


**Supplemental Figure 2** Two independent CoIP repetitions of CLV1-GFP with MAZ-mCherry.

Following the same protocol as described above and shown in Figure 2 A, transiently transformed *N. benthamiana* leaf material (two independent Agrobacterium cultures, different plants, and separate CoIP reactions) harboring the indicated combinations of *XVE*-driven inducible transgenes was deployed to show the capacity of CLV1-GFP to pull down MAZ-mCh. MAZ-mCh alone is not immuno-precipitated via anti-GFP beads (ChromoTek). The Precision Plus Protein™ Dual Color Standard (Bio-Rad) was used for SDS-PAGE (band sizes indicated left).

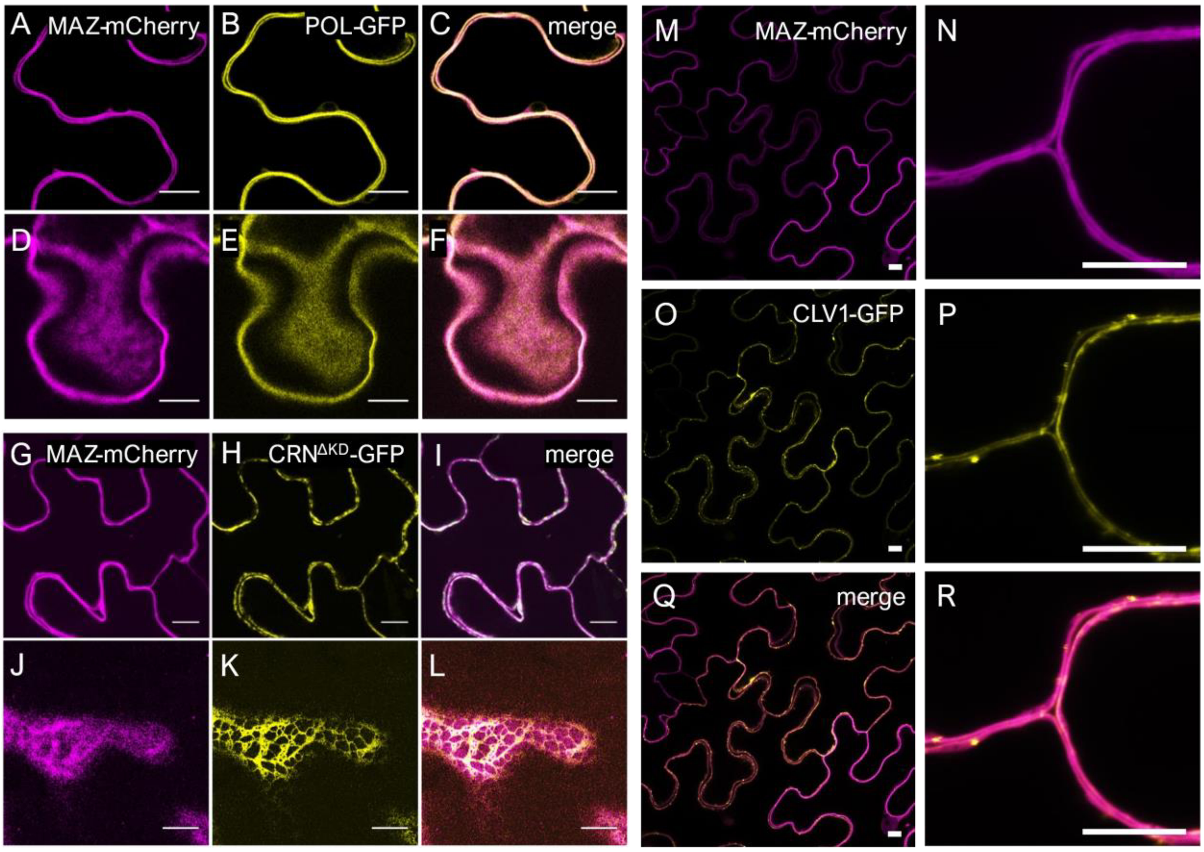


**Supplemental Figure 3** Subcellular localization of MAZ in *N. benthamiana* leaf epidermis cells.

**A – F** Co-expression of *MAZ-mCherry (mCh)* and *POLTERGEIST(POL)-GFP* results in a high degree of local overlap between the two fluorescence signals. POL is known to be localized at the PM (Gagne and Clark, 2010). **G – F** MAZ-mCh does not display notable co-localization with CRN$^{\Delta KD}$-GFP (that is ER-localized in the absence of CLV2). In D – F, and J – L the cell surface is in focus to visualize homogenous distribution of PM-located proteins and network-like structure of ER-associated signals, respectively. **H – R** CLV1-GFP and MAZ-mCh co-localize at the PM, but MAZ is not present in CLV1-GFP vesicles, which probably indicate receptor internalization. All constructs were expressed under the control of the *XVE* promotor system for estradiol induced expression. Scale bars: 10 µm.

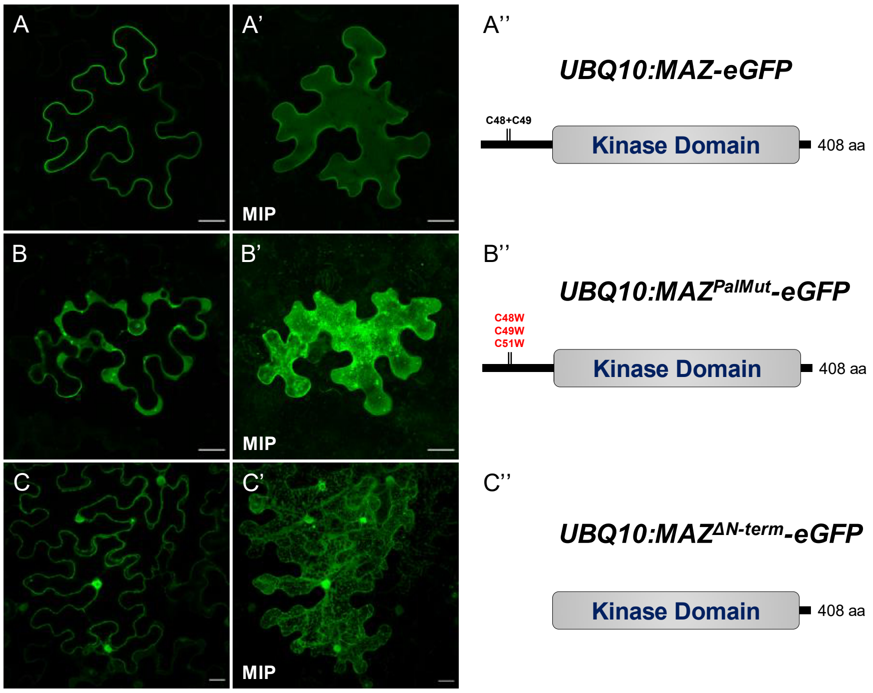


**Supplemental Figure 4** Predicted palmitoylation sites at the N-terminus of MAZ are mandatory for PM-localization.

Transient expression of different *MAZ* variants demonstrate the impact of the N-terminus and the predicted palmitoylation sites C48, C49 (and the adjacent C51) on subcellular localization of MAZ-eGFP. Representative images of *N. benthamiana* leaf epidermis cells expressing the transgenes under the control of the *UBQ10* promotor (**A, B, C**), the corresponding maximum intensity projections (MIP) of z-stacks through the epidermal layer (**A', B', C'**) and schematic representations of the MAZ protein structure in the indicated constructs (**A'', B'', C''**). **A – A''** The full-length MAZ protein (fused to eGFP) is located at the PM. **B – B''** Site directed mutagenesis of the palmitoylation sites causes a shift of protein localization from the PM to the cytoplasm. **C – C''** Deletion of the entire N-terminus (the deletion variant starts directly with the kinase domain) results in protein localization within the cytoplasm, around the ER, and in the nucleus. Sale bars: 25 µm.

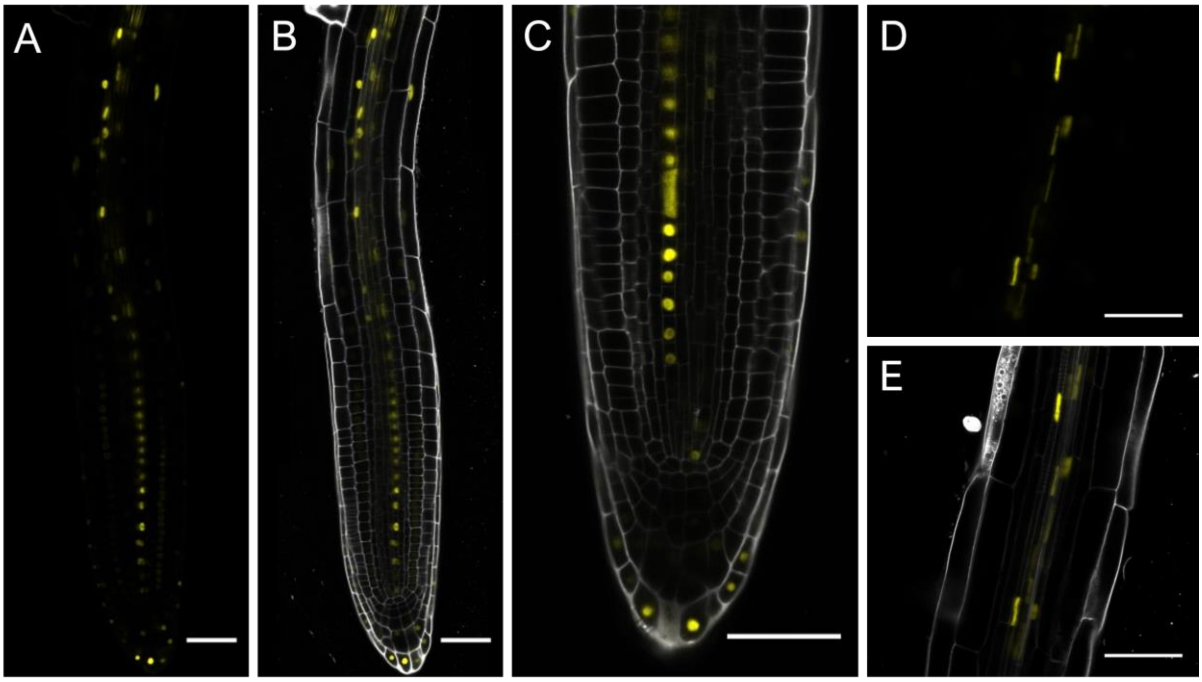


**Supplemental Figure 5** Expression pattern of the transcriptional reporter *MAZ:mVenus-NLS*//Col-0 in the root (3 DAG).

**A, B** The *MAZ* promotor mediates expression predominantly in the vasculature and less pronounced also in other cell files within the meristematic zone of the root. **C** Occasionally observed dispersal of the *per se* nucleus-located fluorescence reporter suggests organelle degradation typical for phloem cells. **D, E** Within the differentiation zone most of the signal is found in the dispersed non-nucleic form, indicating expression within sieve elements. All shown samples are counterstained with PI, merge of mVenus and PI channels in B, C, E. Scalebars: 50 µm.

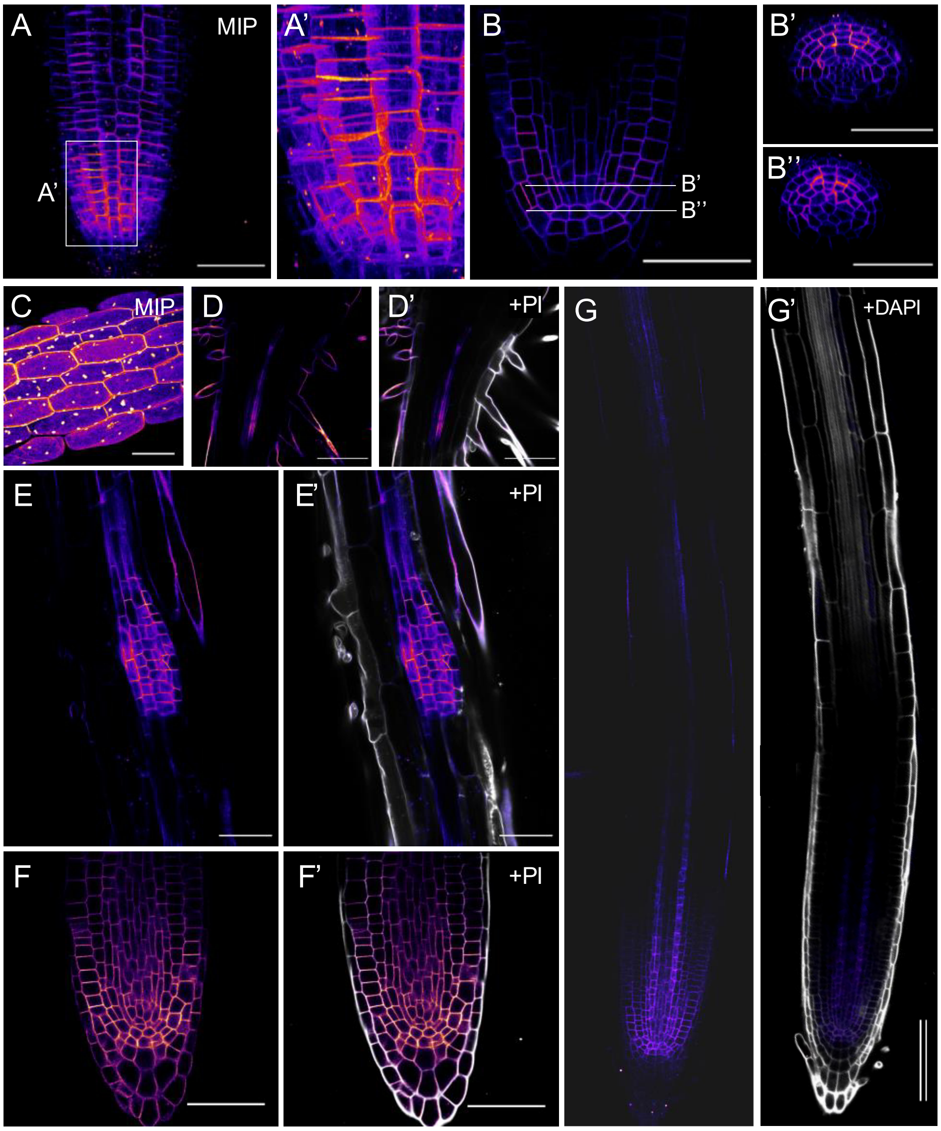


**Supplemental Figure 6** *MAZ* expression by *MAZ* promotor in different translational reporters.

**A – B''** *MAZ-eGFP* expression in the root meristem. The fusion protein is PM-located and found in the QC, initials, and the adjacent cells as different projections of a z-stack through the root tip show, i.e. maximum intensity projection (MIP, A, A'') and YZ cross sections (B', B'') as indicated in B (5 DAG). **C** MIP of a z-stack through the epidermal layer of the hypocotyl of a *MAZ-eGFP* expressing seedling (5 DAG). **D, D'** In the transition region between root and hypocotyl MAZ-eGFP is found in the vasculature and in root hair cells (5 DAG). **E, E'** MAZ-eGFP is present in emerging lateral roots (10 DAG). **F, F'** *MAZ-eGFP* in mature lateral roots resembles its expression in the main root meristem (10 DAG). **G, G'** *MAZ* expression in the root is not dependent on the fluorophore fusion, since MAZ-mCh is found, like MAZ-eGFP, in the vasculature, root hair cells, and the RM (3 DAG). Scalebars: 50 µm (A, D–G), 100 µm (D, D'). Fluorophore signals are visualized with the LUT fire, in D', E', F' merge with PI (grey), in G' merge with DAPI (grey).

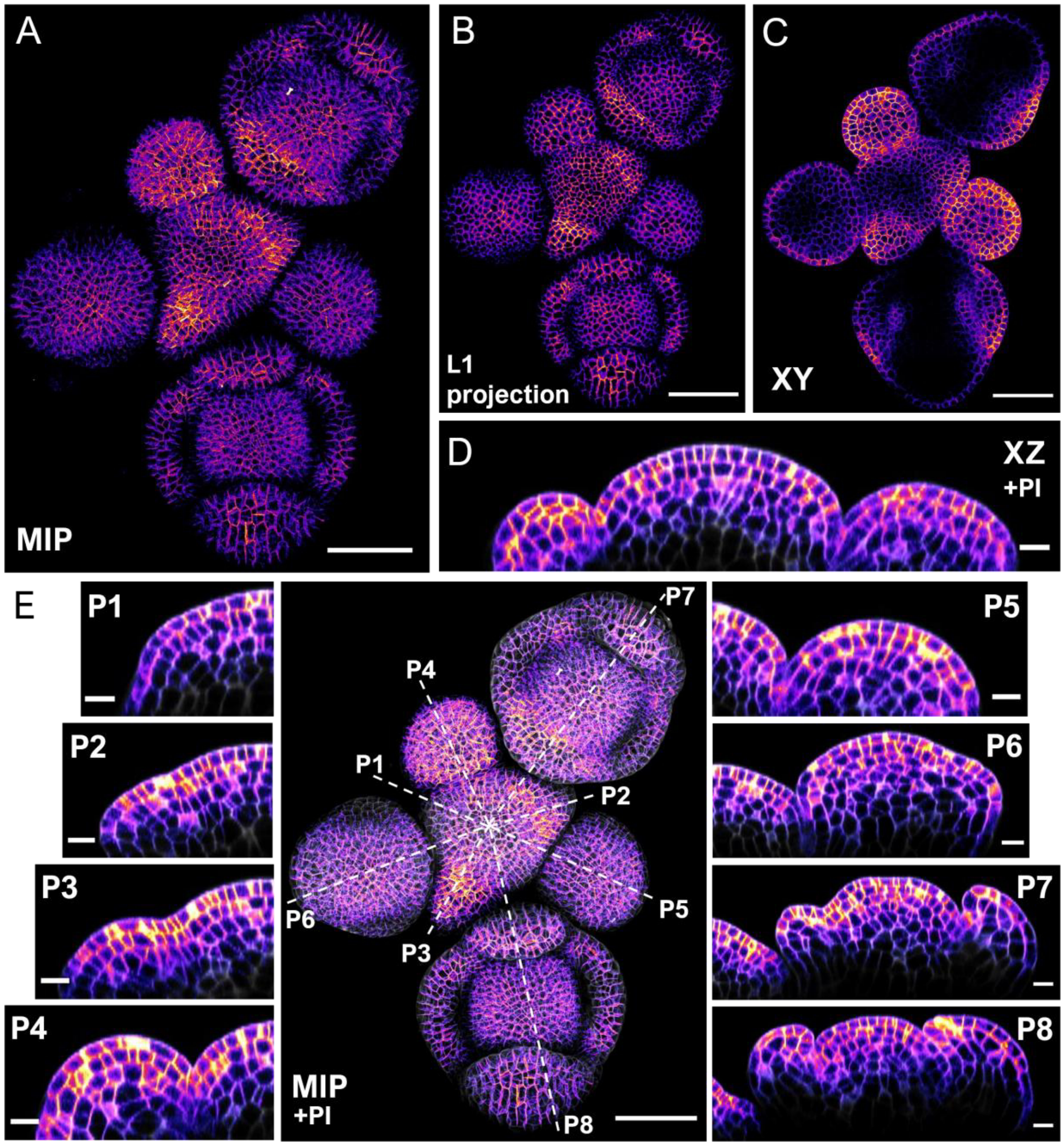


**Supplemental Figure 7** Expression of *MAZ:MAZ-eGFP* in the SAM of 5 weeks old *A. thaliana*.

Various representations of a z-stack through the inflorescence with flower primordia (P) expressing *MAZ-eGFP* driven by the *MAZ* promotor and counterstained with PI (GFP-channel only visualized by LUT fire in A - C; merge with PI in grey displayed in D, E, P1 – P8). **A** The maximum intensity projection (MIP), **B** the L1 projection, **C** a transversal (XY) view, and **D** an orthogonal section through the central region (and P4 + P5), respectively, all indicate a broad *MAZ* expression with areas of higher signal intensity. Those include the boundary regions towards newly formed primordia and towards organs in later stages of development. In general, L1 expression appears elevated with a decreasing intensity gradient towards the L3 layers. **E** MIP as in A but merged with PI channel and cross sections through all primordia (indicated by dotted lines). Scalebars: 50 µm (A, B, C, E), 10 µm (D and primordia (P1 – P8) transversals).

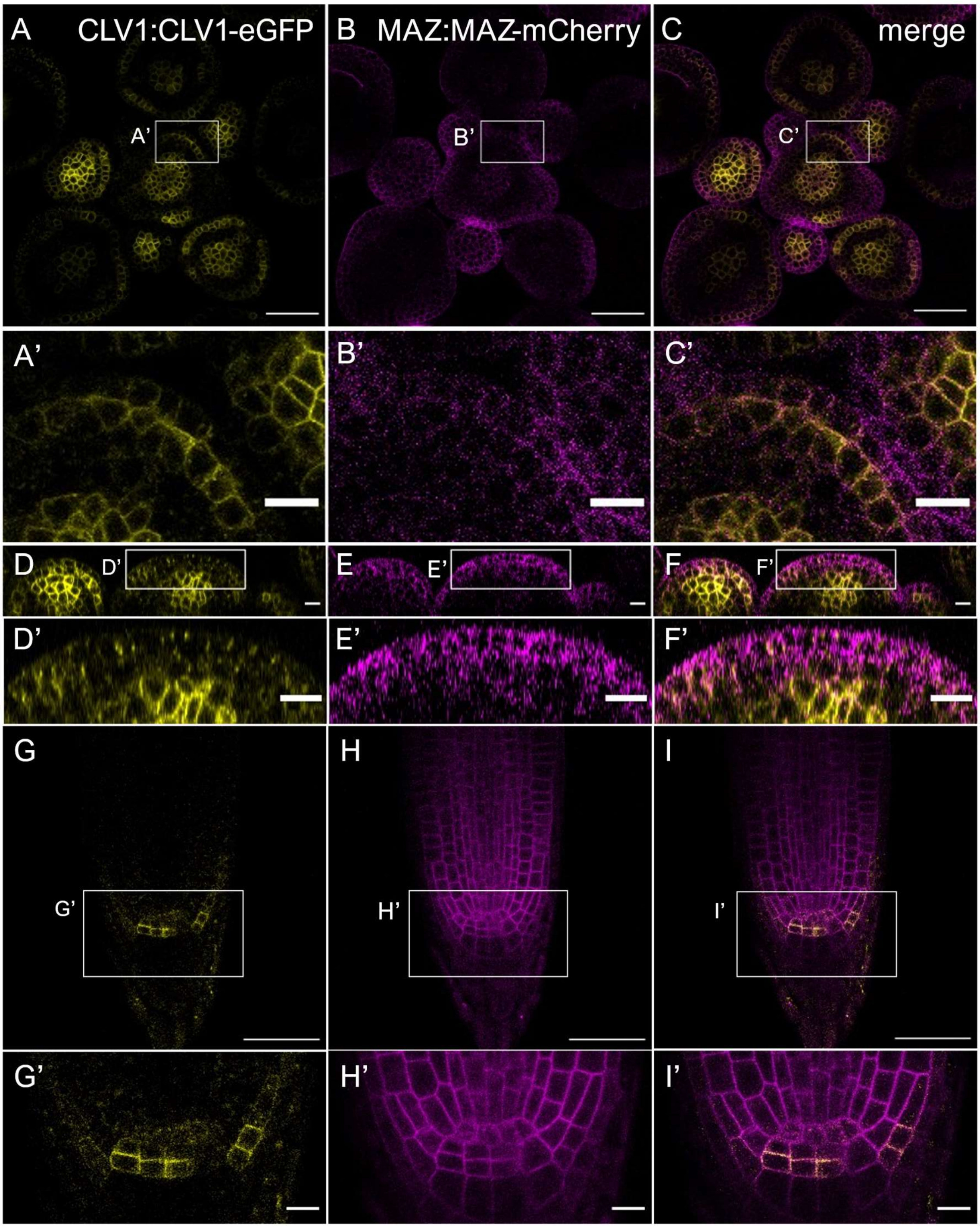


**Supplemental Figure 8** Co-localization of CLV1-eGFP and MAZ-mCherry in shoot and root.

**A – F** Expression pattern of *CLV1* and *MAZ* under the control of the respective endogenous promotors in SAM and flower primordia of 5 weeks old plants. Both, the transversal (XY) cut through the meristem (A – C) and the orthogonal view (XZ, D – F), display CLV1-eGFP in the central zone and in specific areas of the L1, while *MAZ-mCh* is expressed in the entire L1 and only slightly in the central zone. Note the general weak expression of this mCh fusion in the shoot. Nevertheless, the expression pattern is comparable with *MAZ-eGFP* lines (compare Fig. 5). **G – I** In the distal root meristem *CLV1* expression is limited to few cells, like the columella stem cells (5 DAG). Here, CLV1 co-localize with MAZ. However, in the root meristem the *MAZ* expression domain is extended to the complete meristematic zone. A' – I' display the indicated areas in A – I. Scale bars: 50 µm (A – C, G – I), 20 µm (D – F), 10 µm (A' – I').

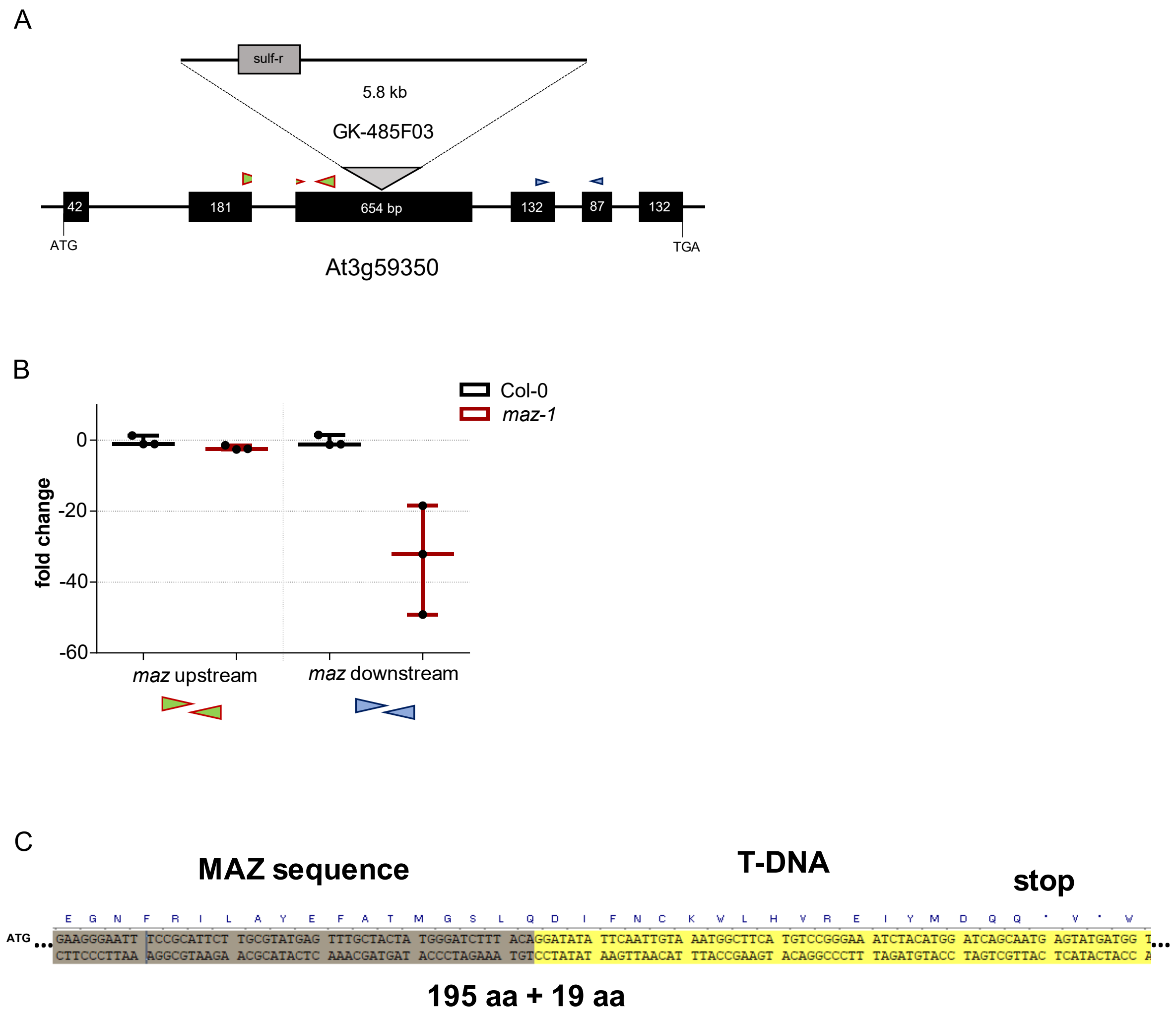


**Supplemental Figure 9** Characterization of the *maz-1* allele GABI-Kat 485F03.

**A** The schematic representation of the genomic region of the *MAZ* gene At3g59350 displays the exon-intron structure as found in the TAIR database and verified by sequencing of cDNA clones (data not shown). Further, the predicted position of the GABI-Kat T-DNA insertion and the binding sites of the two oligo pairs used for quantitative expression analysis are shown. **B** RT-qPCR results from material of Col-0 and *maz-1* (whole seedlings 7 DAG) show a massive decrease of transcripts (up to 50-fold) in the mutant compared to the WT if the oligos bind downstream of the T-DNA insertion. This indicates that the mutant does not express full length MAZ protein anymore. However, there is also a decrease of transcript levels in *maz-1* compared to Col-0 when using an oligo pair upstream of the T-DNA, suggesting instability of the residual mutant *MAZ* mRNA. **C** Transition region from the *MAZ* sequence to the T-DNA insertion start in *maz-1* verified by Sanger sequencing. The insertion event leads to a disruption of the amino acid (aa) sequence of the MAZ protein after 195 aa. The reading frame continuous for 19 triplets of the T-DNA until the first in-frame STOP codon.

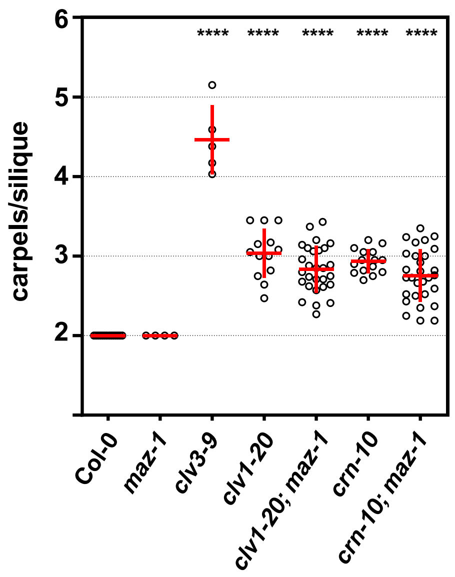


**Supplemental Figure 10** Carpel number of m*az-1* mutants is wild typic in continuous light (CL).

Mean carpel number per silique of indicated genotypes monitored in 6–8 weeks old plants grown under CL. For each genotype 5–30 plants were analyzed by counting the carpels of 15–30 siliques per plant. The average carpel numbers for each plant are plotted. P-values calculated by ANOVA and Dunnett´s post hoc test with **** $p \leq 0.0001$. Sample mean and SD displayed in red.

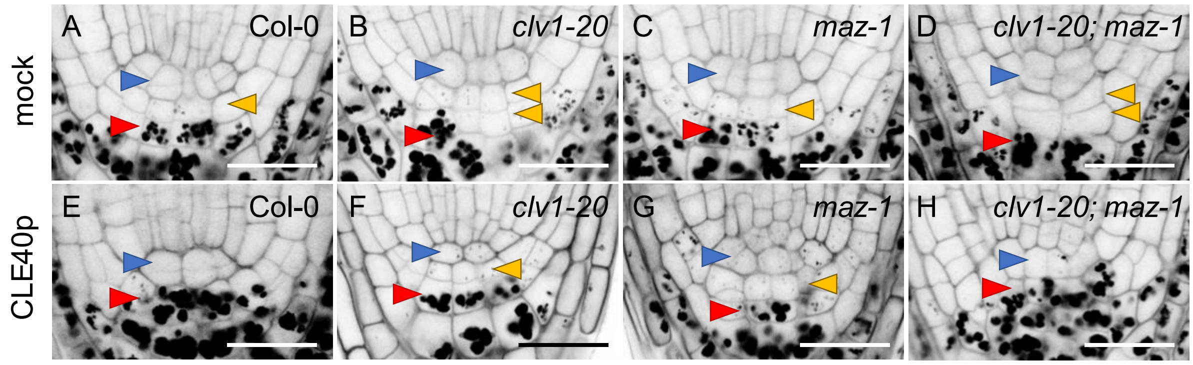


**Supplemental Figure 11** *Maz-1* mutants are partially resistant to CLE40p treatment in terms of columella stem cell (CSC) layer specification.

Representative images of root meristem of mPS-PI stained seedlings (5 DAG) of indicated genotypes, grown on ½ MS plates with mock (A – D) or with CLE40p (200 nM, E – H). Arrow heads mark the QC cells (blue), CSC layers (yellow), and differentiated columella cells (red) with starch granules. Compare Fig. 6 B for quantification.

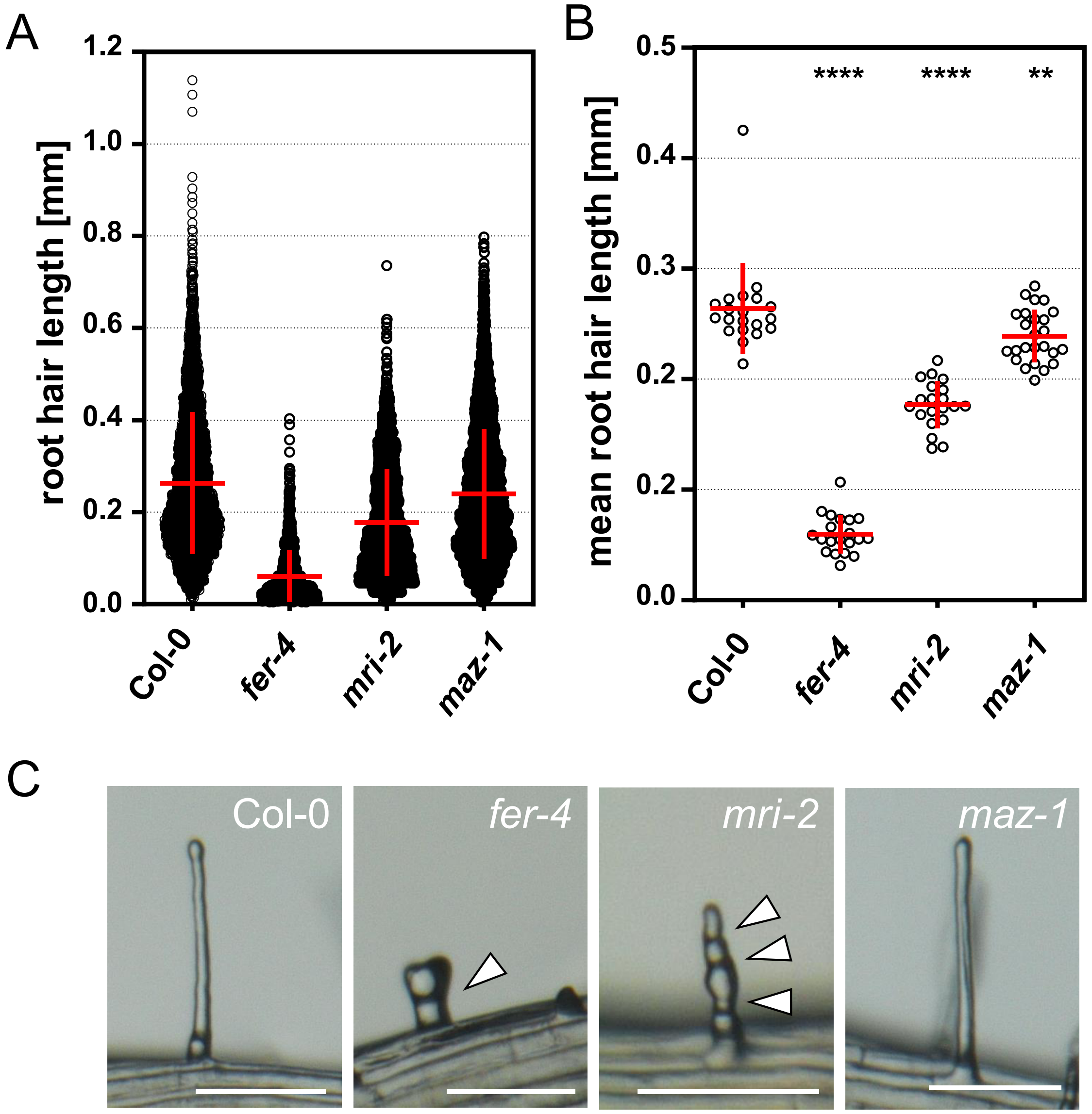


**Supplemental Figure 12** The *maz-1* mutant does not display defects in root hair development.

Seedlings of indicated genotypes were grown on ½ MS agar plates and analyzed regarding root hair development. Root hair length was measured from 20 – 25 roots per genotype (all root hairs in focus on one side of the root) with total n = 3542 (Col-0), 975 (*fer-4*), 2433 (*mri-2*), and 4360 (*maz-1*). Plotting all single measurements (**A**) and the mean values per plant (**B**) revealed that *fer-4* and *mri-2* mutants show reduction of root hair length, while *maz-1* root hair length is comparable to Col-0 samples. P-values in B were calculated by ANOVA and Dunnett´s post hoc test, with ** for $p \leq 0.01$, and **** for $p \leq 0.0001$. **C** Exemplarily selected stereomicroscopy images of root hairs of the analyzed genotypes. Arrow heads show constrictions in the root hairs, probably caused by defective tip growth. This morphology was not observed in *maz-1* mutants. Scale bars: 100 µm.

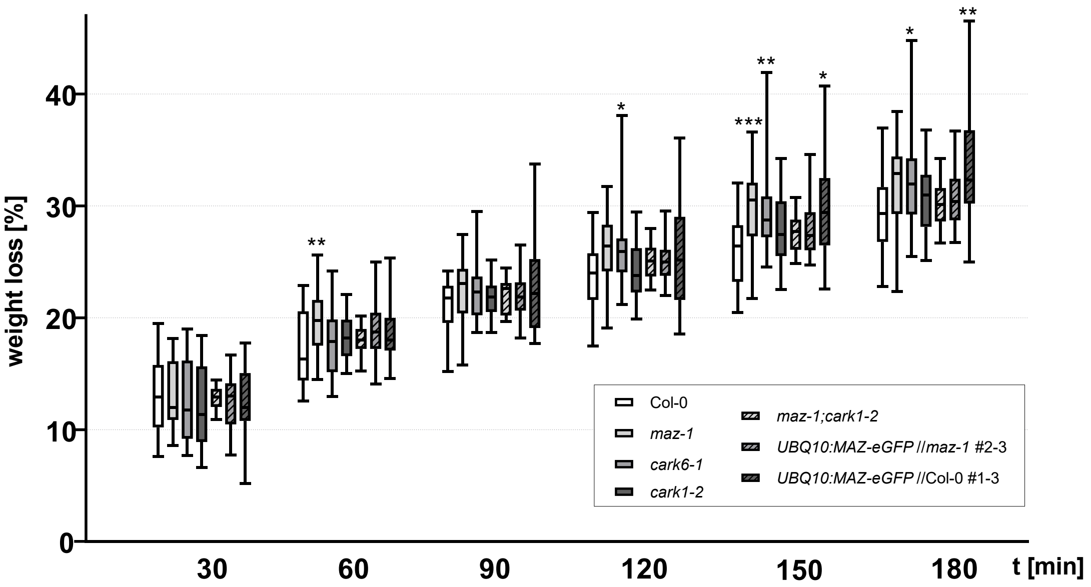


**Supplemental Figure 13** Water loss assay of Arabidopsis leaves to monitor potential differential stomata closure as an indication for a role of Pti1-homologs in ABA signaling.

To estimate water loss through wilting, the reduction of leave fresh weight over time was accessed. The weight of 6 rosette leaves per sample/plant was determined directly after clipping off the leaves at the petioles and again at the indicated time points. The boxplots indicate the distribution of relative weight loss in comparison to the initial measurement for each time point. Alleles of different *pti1-like* mutants, (*maz-1*, *cark6-1*, *cark1-2*, and the double mutant *maz-1;cark1-2*) were included in the assay, as well as a *maz-1* complementing line, and a *MAZ* OE line. For each genotype 24 individual plants were analyzed in 3 independent experiments (in the case of *maz-1;cark1-2* only 16 plants in 2 independent experiments). Plants were cultivated for 4 – 5 weeks under standard LD conditions. The p-values were calculated by ANOVA and Dunnett´s post hoc test in comparison to the Col-0 WT sample for each time point, with * for $p \leq 0.05$, ** for $p \leq 0.01$, and *** for $p \leq 0.001$.

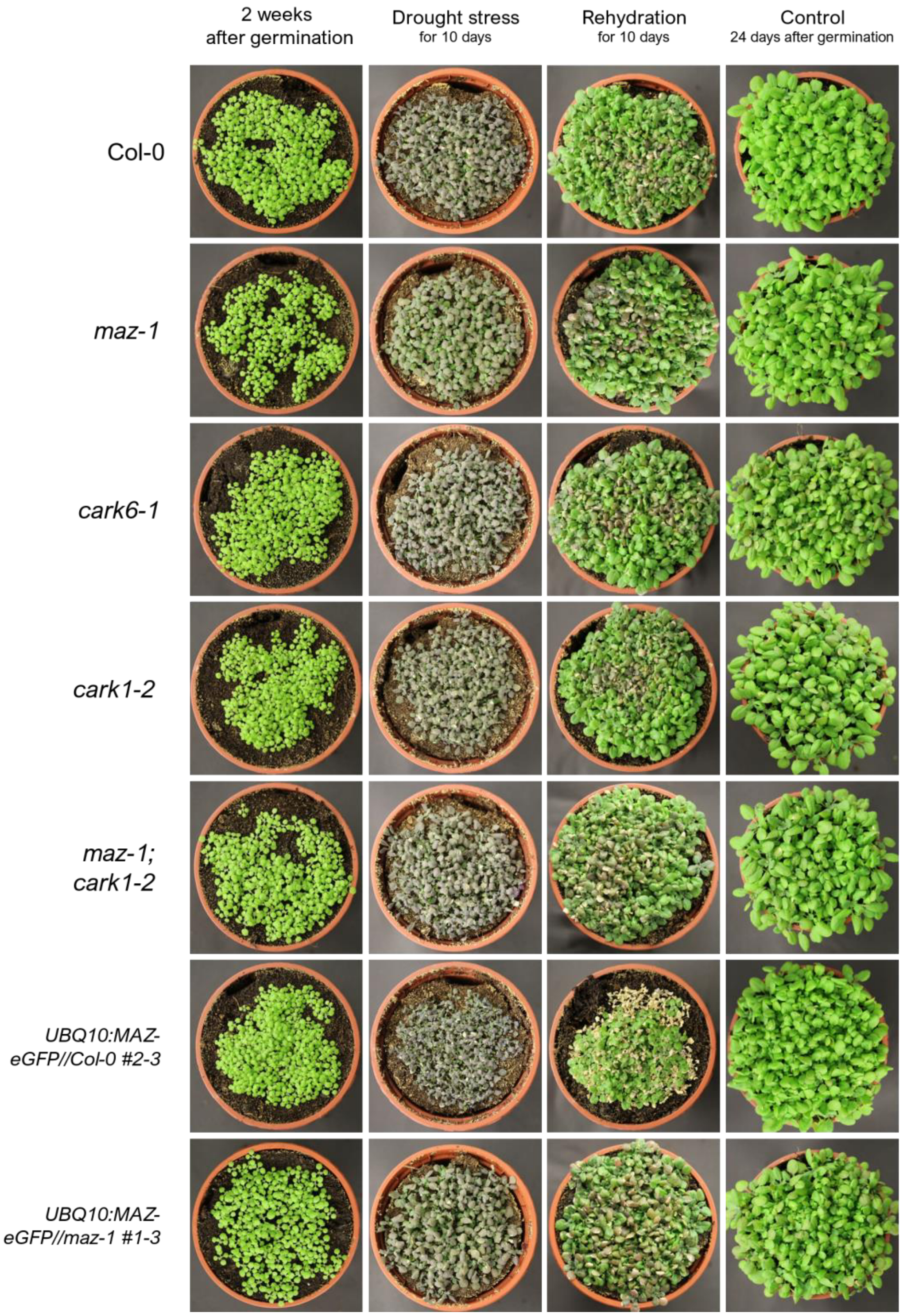


**Supplemental Figure 14** Drought stress assay revealed no clear differences between the analyzed genotypes to cope with water deficiency.

Plants were cultivated under standard LD conditions for 2 weeks (images in column 1). During the following 10 days plants were kept in drought (column 2) und subsequently supplied with water again. Recovery was monitored after 10 days (column 3). The control plants were watered regularly (column 4, 24 days after germination). Representative images from 3 repetitions, all genotypes were randomly distributed over the tray to minimize local effects.

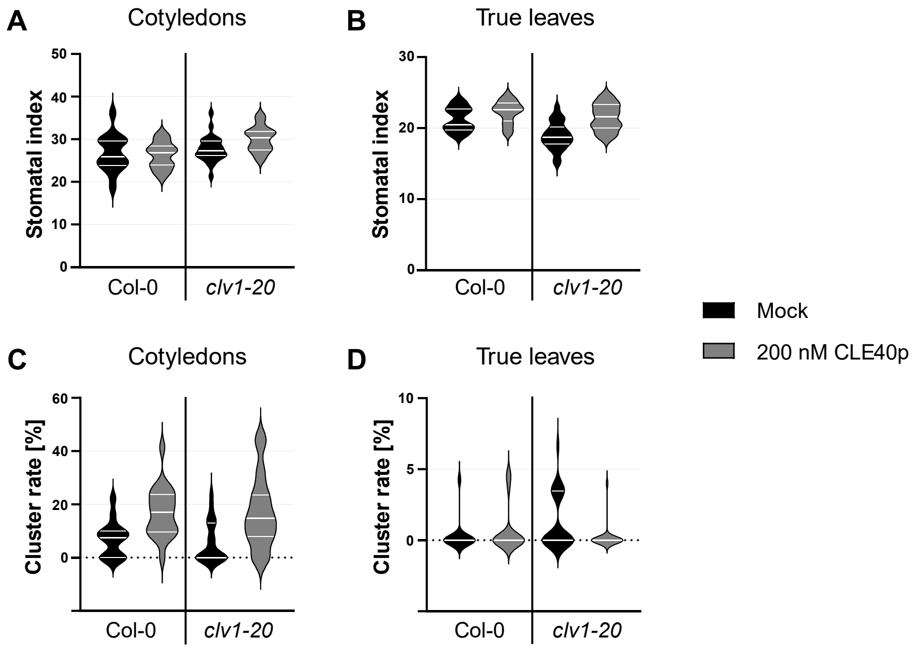


**Supplemental Figure 15** Impact of the *clv1-20* allele and CLE40 peptide treatment on stomata development and clustering in Arabidopsis seedlings.

**A, B** The stomatal index is constant between Col-0 WT and *clv1-20* mutants in cotyledons (A), as well as in true leaves (B). Suppling 200 nM CLE40p to the growth medium (½ MS) does not affect the stomatal index. **C, D** The cluster rate of Col-0 and *clv1-20* is not altered in cotyledons, but CLE40p treatment increases clustering (C). In true leaves *clv1-20* mutants display increased clustering of stomata, which is counteracted by treatment of CLE40p (D). All plants were analysed 14 DAG.

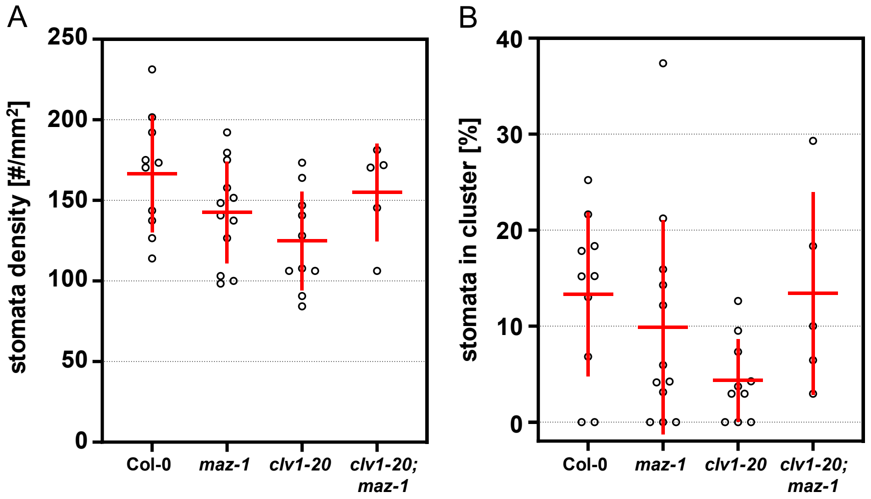


**Supplemental Figure 16** Cotyledons of *clv1-20*, *maz-1*, and the double mutant *clv1-20; maz-1* display no significant changes of stomata density and cluster rate.

**A** Stomatal density and **B** cluster rate were accessed from cotyledons of indicated genotypes. Each plotted data point represents the mean value of 4 areas within one cotyledon of individual plants. Seedlings were grown on ½ MS agar plates. 14 DAG cotyledons were stained with PI to image abaxial epidermis cells. No significant differences in comparison to the Col-0 sample were identified by ANOVA and Dunnett´s post hoc test. Sample mean and SD displayed with red lines.
