## Supplemental Tables 1 - 11 for "The receptor-like cytoplasmic kinase MAZZA and CLAVATA-family receptors interact *in vivo*, together mediating developmental processes in *Arabidopsis thaliana*"

**Supplemental Table 1** Via mass spectroscopy identified proteins in CoIP fraction against CLV1-2xGFP, which are not found in GFP-only control samples.

| Gene ID | Protein name | Subcellular localization | MS/MS hits |
|---|---|---|---|
| At1g75820 | CLAVATA1 (CLV1) | PM | 23 |
| At5g38530 | Tryptophan synthase beta chain (TSBtpye2) | chloroplast, cytosol | 20 |
| At3g56650 | PsbP domain-containing protein 6 (PPD6) | chloroplast | 20 |
| At4g24510 | Protein ECERIFERUM 2 (CER2) | nucleus, ER | 19 |
| At3g10060 | Peptidyl-prolyl cis-trans isomerase (FKBP16-4) | chloroplast | 16 |
| At3g01420 | Alpha-dioxygenase 1 (DOX1) | n/a | 16 |
| At1g16080 | uncharacterized protein | apoplast, ER | 16 |
| At4g11290 | Peroxidase 39 (PER39) | extracellular | 15 |
| At3g06350 | Bifunctional 3-dehydroquinate dehydratase/shikimate dehydrogenase (EMB3004) | chloroplast | 15 |
| At2g46520 | Exportin-2 (CAS) | nucleus | 12 |
| At1g72160 | Patellin-3 (PATL3) | PM, cytoplasm | 11 |
| At2g26250 | 3-ketoacyl-CoA synthase 10 (FDH) | ER | 11 |
| At5g47840 | Adenylate kinase 2 | chloroplast stoma | 10 |
| At5g55730 | Fasciclin-like arabinogalactan protein 1 (FLA1) | PM, apoplast | 10 |
| At4g28440 | uncharacterized protein | cytosol | 10 |
| At4g12470 | pEARLI1-like lipid transfer protein 1 (AZI1) | CW, plasmodesmata, ER | 10 |
| At5g18660 | Divinyl chlorophyllide a 8-vinyl-reductase (DVR) | chloroplast | 9 |
| At5g05730 | Anthranilate synthase alpha subunit 1 (ASA1) | chloroplast | 9 |
| At3g16520 | UDP-glycosyltransferase 88A1 (UGT88A1) | cytosol | 9 |
| At3g59350 | PTI1-like tyrosine-proteine kinase 3 (Pti1-3) | PM-associated | 8 |
| At1g69530 | Expansin-A1 (EXPA1) | CW, extracellular, mitochondria | 7 |
| At3g14100 | Oligouridylate-binding protein 1C (UBP1C) | nucleus | 6 |
| At2g37760 | Aldo-keto reductase family 4 member C8 (AKR4C8) | cytosol | 5 |

**Supplemental Table 2** Mean FRET efficiencies of tested donor-acceptor combinations at the PM of transiently transformed *N. benthamiana* epidermis cells.

| Donor | Acceptor | FRET efficiency [%] ± SD |
|---|---|---|
| CLV1-GFP | MAZ-linker-mCherry | 5.9 ±3.7 |
| CLV1-GFP + 5 µM CLV3 | MAZ-linker-mCherry | 5.5 ±1.8 |
| CLV1-GFP | CRN-KD$^{myr}$-mCherry | 2.0 ±2.8 |
| BAM1-GFP | MAZ-linker-mCherry | 6.5 ±2.3 |
| MAZ-eGFP | MAZ-linker-mCherry | 5.3 ±2.8 |
| MAZ-eGFP | BAM3-mCherry | 4.0 ±2.0 |
| MAZ-eGFP | CIK2-mCherry | 5.7 ±1.9 |
| CRN-GFP (+CLV2) | MAZ-linker-mCherry | 3.0 ±2.3 |
| CLV1-mNeonGreen | MAZ-mCherry | 3.7 ±1.8 |
| CLV1-mNeonGreen | mCherry$^{myr}$ | 0.7 ±1.1 |
| CLV1-mNeonGreen | CARK1-mCherry | 5.2 ±3.2 |
| CLV1-mNeonGreen | Pti1-11-mCherry | 5.4 ±2.2 |
| CLV1-mNeonGreen | CARK6-mCherry | 4.2 ±3.2 |
| CLV1-mNeonGreen | Pti1-2-mCherry | 0.6 ±2.3 |

**Supplemental Table 3** Pti1-like family in *Arabidopsis thaliana* and predicted PM-localization mechanisms.

| Name | TAIR locus | Length [aa] | Myristoylation[1] | Palmitoylation[2] | GPI-anchor[3] |
|---|---|---|---|---|---|
| Pti1-1 | At1g06700 | 361 | no | C6, C7, C9 | no |
| Pti1-2 | At2g30740 | 366 | no | C6, C7 | no |
| MAZZA | At3g59350 | 408 | no | C48, C49 | no |
| Pti1-4 | At2g47060 | 365 | no | C3, C6, C7 | no |
| MARIS | At2g41970 | 365 | no | C3, C4 | no |
| Pti1-6 | At2g30730 | 338 | no | no | no |
| CARK6 | At2g43230 | 440 | no | C79, C80, C82 | no |
| Pti1-8 | At1g48220 | 364 | no | C3, C7 | no |
| Pti1-9 | At3g62220 | 361 | no | C3, C6, C7 | no |
| CARK1 | At3g17410 | 364 | score = 0.6* | C3, C6, C7 | no |
| Pti1-11 | At1g48210 | 363 | no | C3, C7 | no |

[1] predicted by CSS-Palm 4.1
[2] predicted by Myristoylator (ExPASy)
[3] predicted by PredGPI

* indicating medium confidence

**Supplemental Table 4** Chemicals.

| Name | Producer/Source | Product no. | CAS no. |
|---|---|---|---|
| *Acetosyringone (3',5'-Dimethoxy-4'-hydroxy-acetophenone)* | Sigma-Aldrich (Merck) | D134406 | 2478-38-8 |
| *Aniline blue (water soluble)* | Thermo Fisher Scientific | B8563 | 28631-66-5 |
| *BASTA® non-selective herbicide* | Bayer CropScience | 84442615 | N/A |
| β-estradiole | Sigma-Aldrich (Merck) | E8875 | 50-28-2 |
| *Carbenicillin disodium salt* | Carl Roth | 6344.2 | 4800-94-6 |
| *cOmplete*™ Protease Inhibitor Cocktail | Roche | 11697498001 | N/A |
| *DAPI (4',6-diamidino-2-phenylindole)* | N/A | N/A | 28718-90-3 |
| *DL-phosphinothricin (PPT)* | Duchefa Biochemie bv | P0159 | 77182-82-2 |
| *Gentamicin sulfate* | Sigma-Aldrich (Merck) | G1264 | 1405-41-0 |
| *GFP antibody [3H9] (IgG, rat monoclonal)* | ChromoTek | 3h9-20 | N/A |
| *GFP-Trap® magnetic agarose* | ChromoTek | gmt-20 | N/A |
| *Goat anti-mouse IgG Alkaline Phosphatase (AP)* | Sigma-Aldrich (Merck) | AP124A | N/A |
| *Goat anti-rat IgG Alkaline Phospatase (AP)* | Sigma-Aldrich (Merck) | A8438 | N/A |
| *Hygromycin B* | Duchefa Biochemie bv | H0192 | 31282-04-9 |
| *Kanamycin monosulfate* | Duchefa Biochemie bv | K0126.0005 | 25389-94-0 |
| *MES hydrate* | Sigma-Aldrich (Merck) | 10240885 | 1266615-59-1 |
| *Murashige & Skoog Medium (+Gamborg B5 vitamins)* | Duchefa Biochemie bv | M0231.0050 | N/A |
| *NEXT GEL* ® 10 % acrylamide | VWR Life Science | M256 | N/A |
| *Nonident*™ P40 substitute | Sigma-Aldrich (Merck) | 100980769 | 9016-45-9 |
| *Phusion High-Fidelity PCR polymerase* | Thermo Fisher Scientific | F530S | N/A |
| *Propidium iodide* | Thermo Fisher Scientific | P1304MP | 25535-16-4 |
| *RFP antibody [6G6] (IgG, mouse monoclonal)* | ChromoTek | 6g6-20 | N/A |
| *Rifampicin* | TCI | R0079 | 13292-46-1 |
| *Spectinomycin HCl pentahydrate* | Duchefa Biochemie bv | S0188 | 22189-32-8 |
| *SsoAdvanced Univerasal SYBR® Green Supermix* | Bio-Rad | 1725271 | N/A |
| *SuperScript*™ reverse transcriptase | Thermo Fisher Scientific | 18080-093 | N/A |
| *Synthetic CLV3 (RCV[Hyp]SG[Hyp]DPLHHH)* | Peptides & Elephants | costumized | N/A |
| *Synthetic CLE40 (RQV[Hyp]TGSDPLHHK)* | Peptides & Elephants | costumized | N/A |
| *Synthetic CLE45 (RRVRRGSDPIHN)* | Peptides & Elephants | costumized | N/A |
| *Tetracycline* | Sigma-Aldrich (Merck) | 87128 | 60-54-8 |

**Supplemental Table 5** Analyzed *A. thaliana* mutants (Col-0 background).

| Allele | Type | Gene | Reference |
|---|---|---|---|
| *maz-1* | T-DNA | At3g59350 | GABI-Kat 485F03 |
| *clv1-20* | T-DNA | At1g75820 | Durbak and Tax, 2011; SALK 008670 |
| *clv3-9* | EMS | At2g27250 | Rüdiger Simon, 2003; W62Stop |
| *crn-10* | CRISPR/Cas9 | At5g13290 | Nimchuk, 2017 |
| *cark1-2* | T-DNA | At3g17410 | SALK 094451 |
| *cark6-1* | T-DNA | At2g43230 | Wang et al. 2019; SALK 203094 |
| *mri-2* | T-DNA | At2g41970 | Boisson-Dernier et al. 2015; GABI-Kat 820D05 |
| *fer-4* | T-DNA | At3g51550 | Haruta et al. 2014; GABI-Kat 106A06 |

**Supplemental Table 6** Genotyping strategies to verify listed mutant alleles.

| Allele | Method | Details |
|---|---|---|
| *maz-1* | PCR | Fwd 5'-CAGCTCAGTTTGTTAGGCTGGATA-3'<br>Rev (WT) 5'-CAGTATTTTATGGTTAGCGTTTTG-3'<br>Rev (T-DNA) 5'-ATAATAACGCTGCGGACATCTACATTTT-3' (GABI o8474)<br>WT amplicon = 1403 bp; mutant amplicon = 750 bp |
| *clv1-20* | PCR | Fwd 5'-TTTGAATAGTGTGTGACCAAATTTGA-3'<br>Rev (WT) 5'-TCCAATGGTAATTCACCGGTG-3'<br>Rev (T-DNA) 5'-TGGTTCACGTAGTGGGCCATCG-3' (SALK LBa1)<br>WT amplicon = 860 bp; mutant amplicon = 1200 bp |
| *crn-10* | dCAPS | Fwd 5'-GTAGAAGCAGCAATGAAGCAAAGAAGAAGGTG-3'<br>Rev 5'-GTGTAGATGATGTTGAAGTTGTGGATAAGTG-3'<br>subsequent HphI digestion, WT = 128 + 41 bp, mutant = 169 bp |
| *cark1-2* | PCR | Fwd (WT) 5'-TCAACACACTGCTTCACCTTG-3'<br>Fwd (T-DNA) 5'-ATTTTGCCGATTTCGGAAC-3' (SALK LBa1.3)<br>Rev 5'-GAAAAGGTGTTAAAGGGGCAC-3'<br>WT amplicon = 1066 bp; mutant amplicon = 700 bp |
| *cark1-6* | PCR | Fwd (WT) 5'-TGTTCGATTGGATAAACCGAG-3'<br>Fwd (T-DNA) 5'-ATTTTGCCGATTTCGGAAC-3' (SALK LBa1.3)<br>Rev 5'-CCCGGTAAGAAGTTCCAAAAG-3'<br>WT amplicon = 1169 bp; mutant amplicon = 725 bp |
| *mri-2* | PCR | Fwd 5'-GTTCTATTCTTCGACCAAATGG-3'<br>Rev (WT) 5'-CTGCATACTGGTTTGCGGG-3'<br>Rev (T-DNA) 5'-GGGCTACACTGAATTGGTAGCTC-3' (GABI o8760)<br>WT amplicon = 935 bp; mutant amplicon = 880 bp<br>(see Boisson-Dernier et al. 2016) |
| *fer-4* | PCR | Fwd (WT) 5'-GATTACTCTCCAACAGAGAAAATCCT-3'<br>Rev (WT) 5'-CGTATTGCTTTTCGATTTCCTA-3'<br>Fwd (T-DNA) 5'-ACGGTCTCAACGCTACCAAC-3'<br>Rev (T-DNA) 5'-TTTCCCGCCTTCGGTTTA-3'<br>WT amplicon = 1254 bp; mutant amplicon = 570 bp<br>(see Haruta et al. 2014) |

### **Supplemental Table 7** Entry plasmids.

| Name | Description (insert) | Gene | Backbone | Bacterial resistance | Oligos for insert amplification, mutagenesis (or reference) |
|---|---|---|---|---|---|
| *MAZ/pENTR* | genomic region of MAZ (START to one codon before STOP, including Introns) | At3g59350 | *pENTR/D-TOPO®* | Kanamycin | 5'-CACCATGTATCCGATGGATTCTGATTAC-3'<br>5'-GGCTTCCTGGACTGGTACAG-3' |
| *pMAZ/pGGA* | promotor region of MAZ (1.8 kb) | At3g59350 | *pGGA000* | Ampicillin | 5'-TTTGGTCTCAACCTCTATAAATTACAGACATTCAAATAC-3'<br>5'-TTTGGTCTCATGTTAGTGAAAGACGGATCG-3' |
| *MAZ/pGGC* | genomic region of MAZ (START to one codon before STOP, including Introns) | At3g59350 | *pGGC000* | Ampicillin | 5'-TTTGGTCTCAGGCTTAATGTATCCGATGGATTC-3'<br>5'-TTTGGTCTCACTGAGGCTTCCTGGACTG-3' |
| *MAZ* $^{PalMut}$/pGGC | C48W, C49W, C51W in MAZ (site-directed mutagenesis of MAZ/pGGC) | At3g59350 | *pGGC000* | Ampicillin | 5'-GATGCGTAGGTGGTTGTGGTGGGCTTGGCACGTTGAAGAAC-3'<br>5'-GTTCTTCAACGTGCCAAGCCCACCACAACCACCTACGCATC-3' |
| *MAZ* $^{\Delta N\text{-}term}$/pGGC | genomic region of MAZ (from F113, w/o STOP, including Introns) | At3g59350 | *pGGC000* | Ampicillin | 5'-TTTGGTCTCAGGCTCAATGTTTGGATCAAAGTCATTG-3'<br>5'-TTTGGTCTCACTGAGGCTTCCTGGACTG-3' |
| *Pti1-2/pGGC* | genomic region of Pti1-2 (START to one codon before STOP, including Introns) | At2g30740 | *pGGC000* | Ampicillin | 5'-TTTGGTCTCAGGCTTAATGCGTAGGTGGATCTGTTGT-3'<br>5'-TTTGGTCTCACTGAGGATTCCGGTACTGGGGCTGG-3' |
| *CARK1/pGGC* | genomic region of CARK1 (START to one codon before STOP, including Introns) | At3g17410 | *pGGC000* | Ampicillin | 5'-AAACGTCTCAGGCTTGATGGGCTGCTTTGGTT-3' 5'-<br>AAACGTCTCACTGAATACGGGTTCCTGTGTGG-3' |
| *CARK6/pGGC* | genomic region of CARK6 (START to one codon before STOP, including Introns) | At2g43230 | *pGGC000* | Ampicillin | 5'-TTTGGTCTCAGGCTATGGATCGTGATTTTCATCG-3'<br>5'-TTTGGTCTCACTGAAGGTTGAGGTGTTGG-3' |
| *Pti1-6/pGGC* | genomic region of Pti1-6 (START to one codon before STOP, including Introns) | At1g48210 | *pGGC000* | Ampicillin | 5'-TTTGGTCTCAGGCTTAATGTTTTTTTTTCTTGTCATCACAT-3'<br>5'-TTTGGTCTCACTGAGAATTGCGGTATTGAGCCTGT-3' |
| *BAM3/pENTR* | BAM3 coding region (with introns) | At4g20270 | *pENTR/D-TOPO®* | Kanamycin | 5'-CACCATGGCAGACAAGATCTTCAC-3'<br>5'-GAAAGTATTAGGCTGTTTAGCC-3' |
| *CIK2/pENTR* | CIK2 coding region | At2g23950 | *pENTR/D-TOPO®* | Kanamycin | Pauline Anne (Christian Hardtke Lab) |
| *POL/pENTR* | POLTERGEIST coding region | At2g46920 | *pENTR/D-TOPO®* | Kanamycin | Frederic Boyer |
| *BAM1/pENTR* | BAM1 coding region | At5g65700 | *pENTR/D-TOPO®* | Kanamycin | Marc Somssich |
| *myr-CRN-KD/pENTR* | KD of CRN with myristoylation motif | At5g13290 | *pENTR/D-TOPO®* | Kanamycin | Marc Somssich |
| *linker-eGFP/pGGD* | 11 aa linker fused upstream to eGFP | - | *pGGD000* | Ampicillin | Grégoire Denay (pGD165) |
| *mVenus/pGGC* | mVenus w/o STOP codon | - | *pGGC00* | Ampicillin | Rebecca Burkart (pRD43) |
| *mCherry/pGGD* | mCherry with STOP codon | - | *pGGD000* | Ampicillin | Rebecca Burkart (pRD53) |

### **Supplemental Table 8** Plasmids used for transient gene expression in *N. benthamiana*.

| Name | Expression cassette | Cloning strategy | Details | Bacterial resistance |
|---|---|---|---|---|
| *BAM1/pAB118* | XVE<<LexA-min35S:BAM1-mCherry | LR reaction | *BAM1/pENTR + pAB118* | Spectinomycin |
| *BAM3/pAB118* | XVE<<LexA-min35S:BAM3-mCherry | LR reaction | *BAM3/pENTR + pAB118* | Spectinomycin |
| *CIK2/pAB118* | XVE<<LexA-min35S:CIK2-mCherry | LR reaction | *CIK2/pENTR + pAB118* | Spectinomycin |
| *CLV1/pAB117* | XVE<<LexA-min35S:CLV1-GFP | LR reaction | *Bleckmann et al. 2010* | Spectinomycin |
| *CLV1/pAB118* | XVE<<LexA-min35S:CLV1-mCherry | LR reaction | *Bleckmann et al. 2010* | Spectinomycin |
| *CLV2 (untagged)* | XVE<<LexA-min35S:CLV2 | LR reaction | *Bleckmann et al. 2010* | Spectinomycin |
| *CRN/pAB117* | XVE<<LexA-min35S:CRN-GFP | LR reaction | *Bleckmann et al. 2010* | Spectinomycin |
| *CRN* $^{\Delta KD}$/pAB117 | XVE<<LexA-min35S:CRN $^{\Delta KD}$-GFP | LR reaction | *Bleckmann et al. 2010* | Spectinomycin |
| *MAZ/pAB118* | XVE<<LexA-min35S:MAZ-mCherry | LR reaction | *MAZ/pENTR + pAB118* | Spectinomycin |
| *myr-CRN-KD/pAB118* | XVE<<LexA-min35S:myr-CRN-KD-mCherry | LR reaction | *myr-CRN-KD/pENTR + pAB118* | Spectinomycin |
| *POL/pAB117* | XVE<<LexA-min35S:POL-GFP | LR reaction | *POL/pENTR + pAB117* | Spectinomycin |
| *UBQ10:MAZ-eGFP/pGGZ001* | pUBQ10:Ω-MAZ-eGFP:tUBQ10<<HygR | GreenGate reaction | *pGGA006, pGGB002, MAZ/pGGC, linker-eGFP/pGGD, pGGE009, pGGF005 + pGGZ001* | Spectinomycin |
| *UBQ10:MAZ-mCherry/pGGZ001* | pUBQ10:Ω-MAZ-mCherry:tUBQ10<<HygR | GreenGate reaction | *pGGA006, pGGB002, MAZ/pGGC, mCherry/pGGD, pGGE009, pGGF005 + pGGZ001* | Spectinomycin |
| *UBQ10:Pti1-2-mCherry/pGGZ001* | pUBQ10:Ω-Pti1-2-mCherry:tUBQ10 | GreenGate reaction | *pGGA006, pGGB002, Pti1-2/pGGC, mCherry/pGGD, pGGE009, dummy/pGGF + pGGZ001* | Spectinomycin |
| *UBQ10:CARK1-mCherry/pGGZ001* | pUBQ10:Ω-CARK1-mCherry:tUBQ10 | GreenGate reaction | *pGGA006, pGGB002, CARK1/pGGC, mCherry/pGGD, pGGE009, dummy/pGGF + pGGZ001* | Spectinomycin |
| *UBQ10:CARK6-mCherry/pGGZ001* | pUBQ10:Ω-CARK6-eGFP:tUBQ10 | GreenGate reaction | *pGGA006, pGGB002, CARK6/pGGC, mCherry/pGGD, pGGE009, dummy/pGGF + pGGZ001* | Spectinomycin |
| *UBQ10:Pti1-6-mCherry/pGGZ001* | pUBQ10:Ω-Pti1-6-mCherry:tUBQ10 | GreenGate reaction | *pGGA006, pGGB002, Pti1-6/pGGC, mCherry/pGGD, pGGE009, dummy/pGGF + pGGZ001* | Spectinomycin |
| *UBQ10:CLV1-mNeonGreen/pGGZ001* | pUBQ10:Ω-CLV1-mNeonGreen:tUBQ10<<HygR | GreenGate reaction | *Grégoire Denay (pGD354)* | Spectinomycin |
| *UBQ10:myr-mCherry/pGGZ001* | pUBQ10:Ω-myr-mCherry:tUBQ10<<HygR | GreenGate reaction | *Grégoire Denay (pGD321)* | Spectinomycin |

**Supplemental Table 9** Plasmids used for stabile transformation of *A. thaliana*.

| Name | Promotor | N-tag | CDS | C-tag | Terminator | Plant Resistance | Backbone |
|---|---|---|---|---|---|---|---|
| *MAZ:MAZ-eGFP+BastaR/pGGZ001* | pMAZ/pGGA | pGGB002 | *MAZ/pGGC* | linker-eGFP/pGGD | pGGE009 | pGGF008 | pGGZ001 |
| *MAZ:MAZ-mCherry+HygR/pGGZ001* | pMAZ/pGGA | pGGB002 | *MAZ/pGGC* | mCherry/pGGD | pGGE009 | pGGF005 | pGGZ001 |
| *UBQ10:MAZ-eGFP+BastaR/pGGZ001* | pGGA006 | pGGB002 | *MAZ/pGGC* | linker-eGFP/pGGD | pGGE009 | pGGF008 | pGGZ001 |
| *MAZ:MAZ$^{PalMut}$-eGFP+BastaR/pGGZ001* | pMAZ/pGGA | pGGB002 | *MAZ*$^{PalMut}$/pGGC | linker-eGFP/pGGD | pGGE009 | pGGF008 | pGGZ001 |
| *UBQ10:MAZ$^{PalMut}$-eGFP+BastaR/pGGZ001* | pGGA006 | pGGB002 | *MAZ*$^{PalMut}$/pGGC | linker-eGFP/pGGD | pGGE009 | pGGF008 | pGGZ001 |
| *MAZ:mVenus-NLS+BastaR/pGGZ001* | pMAZ/pGGA | pGGB002 | mVenus/pGGC | pGGD007 | pGGE009 | pGGF008 | pGGZ001 |

**Supplemental Table 9** Transgenic *A. thaliana* lines applied and generated in this study.

| Name | Transgene description | Reference |
|---|---|---|
| *CLV1:CLV1-2xGFP//clv1-11* | pCLV1:CLV1-2xmGFP/pBJ36 (HygR) | Nimchuk *et al.* 2011 |
| *MAZ:MAZ-eGFP+BastaR//* Col-0 | pMAZ:Ω-MAZ-eGFP:tUBQ10<<pNOS:BastaR:tNOS | this study |
| *MAZ:MAZ-mCh+HygR//* Col-0 | pMAZ:Ω-MAZ-mCherry:tUBQ10<<pUBQ10:HygR:tOCS | this study |
| *MAZ:MAZ$^{PalMut}$-eGFP+BastaR//maz-1* | pMAZ:Ω-MAZ$^{PalMut}$*-eGFP:tUBQ10<<pNOS:BastaR:tNOS* | this study |
| *MAZ:mVenus-NLS+BastaR//* Col-0 | pMAZ:Ω-mVenus-NLS:tUBQ10<<pNOS:BastaR:tNOS | this study |
| *UBQ10:MAZ-eGFP+BastaR//* Col-0 | pUBQ10:Ω-MAZ-eGFP:tUBQ10<<pNOS:BastaR:tNOS | this study |
| *UBQ10:MAZ-eGFP+BastaR//maz-1* | pUBQ10:Ω-MAZ-eGFP:tUBQ10<<pNOS:BastaR:tNOS | this study |
| *UBQ10:MAZ$^{PalMut}$-eGFP+BastaR//* Col-0 | pUBQ10:Ω-MAZ$^{PalMut}$*-eGFP:tUBQ10<<pNOS:BastaR:tNOS* | this study |

**Supplemental Table 11** Oligonucleotides for RT-qPCR.

| Gene | Forward | Revers |
|---|---|---|
| *MAZ* (upstream) | CACAATGATTTTGGGGCATCAC | GCGACAGCCTTTCCATCTTTC |
| *MAZ* (downstream) | TGGTAGGAAACCCGTCGAT | TGGATCAACACATTGCTTCAC |
| AT2G28390 | GGATTTTCAGCTACTCTTCAAGCTA | TCCTGCCTTGACTAAGTTGACA |
| AT4G34270 | GCTCATGGTTCCTCCTCTTG | TCTTCGCCAAACCTATAATGC |
| AT4G26410 | CGTCCACAAAGCTGAATGTG | CGAAGTCATGGAAGCCACTT |
