## Appendix for "The receptor-like cytoplasmic kinase MAZZA and CLAVATA-family receptors interact *in vivo*, together mediating developmental processes in *Arabidopsis thaliana*"

**Appendix** Sequence alignment with ClustalΩ of Arabidopsis Pti1-like family and SlPti1.

```
AtPti1-6      1 ------------------------------------------------------------
AtPti1-1      1 ------------------------------------------------------------
AtPti1-2      1 ------------------------------------------------------------
AtMAZZA       1 MYPMDSDYHRRGL----------------------------------VANDRSPAQFVRL
AtCARK6       1 ---MDRDFHRRGQVVNQDQRATNSNVFTKFENTYLQITAHLCVLVKTQANDRTQSNFVRL
AtMARIS       1 ------------------------------------------------------------
AtPti1-4      1 ------------------------------------------------------------
AtPti1-9      1 ------------------------------------------------------------
AtPti1-8      1 ------------------------------------------------------------
SlPti1        1 ------------------------------------------------------------
AtCARK1       1 ------------------------------------------------------------
AtPti1-11     1 ------------------------------------------------------------

AtPti1-6      1 -------------------------------------------MFFFL-VITYVANQKNQ
AtPti1-1      1 ----------------MRKWICCTCQIED---SNE------EQQLKSS-QQQSDANHKNS
AtPti1-2      1 ----------------MRRWICCGDKKGDSDLSNE------EVHLKSP-WQNSEANQKNQ
AtMAZZA      27 DKPRAVDDLYIGKREKMRRWLCCACHVEEPYHSSE------NEHLRS-PKHHNDFGHHTR
AtCARK6      58 DKPRAVDDLDIGKRGKMRRWLCCSCRVQESYPSAE------NNRLKTPPTRHYDYGRNNK
AtMARIS       1 -------------------MFCCGGADEEPAGPPANQYAAPPNKAGNPNFGGGNRG-EP-
AtPti1-4      1 ----------------MSCFGCCGEDDDMH---KTADYGGRHNQAKHFPPGNDARH-HQ-
AtPti1-9      1 ----------------MSCFGCCREDD-LP---GANDYGGHNMTK---QSGGNDGR-RN-
AtPti1-8      1 ----------------MSCFGWCG-SEDVR---NPADTGPSQAH----NSIGYNGR-HH-
SlPti1        1 ----------------MSCFSCCD-DDDMH---RATDNGPFMAH----NSAGNNGG-QR-
AtCARK1       1 ----------------MGCFGCCGGGEDFR---RVSETGPKPVH----NTGGYNGG-HH-
AtPti1-11     1 ----------------MSCFGWCG-SEDFR---NATDTGPRPAH----NPAGYNGG-HY-

AtPti1-6     17 KPQDLAKP---KEILPIIVPSLSVDEVNEQTDNFGPNSLIGEGSYGRVYYATLNDGKAVA
AtPti1-1     35 KPAPVAKHEVKKEALPIEVPPLSLDEVKEKTENFGSKALIGEGSYGRVYYATLNDGVAVA
AtPti1-2     38 KPQAVVKPEAQKEALPIEVPPLSVDEVKEKTDNFGSKSLIGEGSYGRVYYATLNDGKAVA
AtMAZZA      80 KPQAAVKPDALKEPPSIDVPALSLDELKEKTDNFGSKSLIGEGSYGRAYYATLKDGKAVA
AtCARK6     112 KTPAPVKPPVLKEPPPIDVPAMSLVELKEKTQNFGSKALIGEGSYGRVYYANFNDGKAVA
AtMARIS      40 RNPNAPRSGAPAKVLPIEIPSVALDELNRMAGNFGNKALIGEGSYGRVFCGKFKG-EAVA
AtPti1-4     40 ASETAQKGPPVVKLQPIEVPIIPFSELKEATDDFGSNSLIGEGSYGRVYYGVLNNDLPSA
AtPti1-9     36 GSETAQKGAQSVKVQPIEVAAILADELIEATNDFGTNSLIGEGSYARVYHGVLKNGQRAA
AtPti1-8     35 QRADPPMNQPVVNMQPIAVPAIPVDELEDITENFSSEVLVGKGSYGRVFYGVLKSGKEAA
SlPti1       35 ATESAQRETQTVNIQPIAVPSIAVDELKDITDNFGSKALIGEGSYGRVYHGVLKSGRAAA
AtCARK1      36 QRADPPKNLPVIQMQPISVAAIPADELRDITDNYGSKSLIGEGSYGRVFYGILKSGKAAA
AtPti1-11    35 QRADPPMNQPVIPMQPISVPAIPVDELRDITDNYGSKTLIGEGSYGRVFYGVLKSGGAAA

AtPti1-6     74 LKKLDLAPEDETNTEFLSQVSMVSRLKHENLIQLVGYCVDENLRVLAYEFATMGSLHDIL
AtPti1-1     95 LKKLDVAPEAETDTEFLSQVSMVSRLKHENLIQLLGFCVDGNLRVLAYEFATMGSLHDIL
AtPti1-2     98 LKKLDVAPEAETNTEFLNQVSMVSRLKHENLIQLVGYCVDENLRVLAYEFATMGSLHDIL
AtMAZZA     140 VKKLDNAAEPESNVEFLTQVSRVSKLKHDNFVELFGYCVEGNFRILAYEFATMGSLHDIL
AtCARK6     172 VKKLDNASEPETNVEFLTQVSKVSRLKSDNFVQLLGYCVEGNLRVLAYEFATMRSLHDIL
AtMARIS      99 IKKLDASSSEEPDSDFTSQLSVVSRLKHDHFVELLGYCLEANNRILIYQFATKGSLHDVL
AtPti1-4    100 IKKLD--SNKQPDNEFLAQVSMVSRLKHDNFVQLLGYCVDGNSRILSYEFANNGSLHDIL
AtPti1-9     96 IKKLD--SNKQPNEEFLAQVSMVSRLKHVNFVELLGYSVDGNSRILVFEFAQNGSLHDIL
AtPti1-8     95 IKKLY--PTKQPDQEFLSQVSMVSRLHHENVVALMAYCVDGPLRVLAYEFATYGTLHDVL
SlPti1       95 IKKLD--SSKQPDREFLAQVSMVSRLKDENVVELLGYCVDGGFRVLAYEYAPNGSLHDIL
AtCARK1      96 IKKLD--SSKQPDQEFLAQVSMVSRLRQENVVALLGYCVDGPLRVLAYEYAPNGSLHDIL
AtPti1-11    95 IKKLD--SSKQPDQEFLSQISMVSRLRHDNVTALMGYCVDGPLRVLAYEFAPKGSLHDTL

AtPti1-6    134 HGRKGVQDALPGPTLDWITRVKIAVEAARGLEYLHEKVQPQVIHRDIRSSNILLFDDYQA
AtPti1-1    155 HGRKGVQGAQPGPTLDWITRVKIAVEAARGLEYLHEKSQPPVIHRDIRSSNVLLFEDYKA
AtPti1-2    158 HGRKGVQGAQPGPTLDWLTRVKIAVEAARGLEYLHEKVQPPVIHRDIRSSNVLLFEDYQA
AtMAZZA     200 HGRKGVQGAQPGPTLDWIQRVRIAVDAARGLEYLHEKVQPAVIHRDIRSSNVLLFEDFKA
AtCARK6     232 HGRKGVQGAQPGPTLEWMQRVRVAVDAAKGLEYLHEKVQPAVIHRDIRSSNVLIFEDFKA
AtMARIS     159 HGRKGVQGAEPGPVLNWNQRVKIAYGAAKGLEFLHEKVQPPIVHRDVRSSNVLLFDDFVA
```

```
AtPti1-4    158 HGRKGVKGAQPGPVLSWYQRVKIAVGAARGLEYLHEKANPHIIHRDIKSSNVLLFEDDVA
AtPti1-9    154 HGRKGVKGAKPGPLLSWHQRVKIAVGAARGLEYLHEKANPHVIHRDIKSSNVLIFDNDVA
AtPti1-8    153 HGQTGVIGALQGPVMTWQRRVKIALGAARGLEYLHKKVNPQVIHRDIKASNILLFDDDIA
SlPti1      153 HGRKGVKGAQPGPVLSWAQRVKIAVGAAKGLEYLHEKAQPHIIHRDIKSSNILLFDDDVA
AtCARK1     154 HGRKGVKGAQPGPVLSWHQRVKIAVGAARGLEYLHEKANPHVIHRDIKSSNVLLFDDDVA
AtPti1-11   153 HGKKGAKGALRGPVMTWQQRVKIAVGAARGLEYLHEKVSPQVIHRDIKSSNVLLFDDDVA

AtPti1-6    194 KIADFNLSNQSPDNAARLQSTRV-LGSFGYYSPEYAMTGELTHKSDVYGFGVVLLELLTG
AtPti1-1    215 KIADFNLSNQAPDNAARLHSTRV-LGTFGYHAPEYAMTGQLTQKSDVYSFGVVLLELLTG
AtPti1-2    218 KVADFNLSNQAPDNAARLHSTRV-LGTFGYHAPEYAMTGQLTQKSDVYSFGVVLLELLTG
AtMAZZA     260 KIADFNLSNQSPDMAARLHSTRV-LGTFGYHAPEYAMTGQLTQKSDVYSFGVVLLELLTG
AtCARK6     292 KIADFNLSNQAPDMAARLHSTRV-LGTFGYHAPEYAMTGQLTQKSDVYSFGVVLLELLTG
AtMARIS     219 KMADFNLTNASSDTAARLHSTRV-LGTFGYHAPEYAMTGQITQKSDVYSFGVVLLELLTG
AtPti1-4    218 KIADFDLSNQAPDMAARLHSTRV-LGTFGYHAPEYAMTGQLNAKSDVYSFGVVLLELLTG
AtPti1-9    214 KIADFDLSNQAPDMAARLHSTRV-LGTFGYHAPEYAMTGQLSAKSDVYSFGVVLLELLTG
AtPti1-8    213 KIGDFDLYDQAPNMAGRLHSCRMALGASRSHCPEHAMTGILTTKSDVYSFGVVLLELLTG
SlPti1      213 KIADFDLSNQAPDMAARLHSTRV-LGTFGYHAPEYAMTGQLSSKSDVYSFGVVLLELLTG
AtCARK1     214 KIADFDLSNQAPDMAARLHSTRV-LGTFGYHAPEYAMTGTLSTKSDVYSFGVVLLELLTG
AtPti1-11   213 KIGDFDLSDQAPDMAARLHSTRV-LGTFGYHAPEYAMTGTLSSKSDVYSFGVVLLELLTG

AtPti1-6    253 RKPVDHTMPRGQQSLVTWATPKLSEDTVEECVDPKLKGEYSPKSVAK-------------
AtPti1-1    274 RKPVDHTMPRGQQSLVTWATPRLSEDKVKQCIDPKLKADYPPKAVAK-------------
AtPti1-2    277 RKPVDHTMPRGQQSLVTWATPRLSEDKVKQCVDPKLKGEYPPKSVAK-------------
AtMAZZA     319 RKPVDHTMPRGQQSLVTWATPRLSEDKVKQCVDPKLKGEYPPKAVAK-------------
AtCARK6     351 RKPVDHTMPRGQQSLVTWATPRLSEDKVKQCVDPKLKGEYPPKAVAK-------------
AtMARIS     278 RKPVDHTMPKGQQSLVTWATPRLSEDKVKQCIDPKLNNDFPPKAVAK-------------
AtPti1-4    277 RKPVDHRLPRGQQSLVTWATPKLSEDKVKQCVDARLGGDYPPKAVAKVRNQTFHNLRLCL
AtPti1-9    273 RKPVDHTLPRGQQSLVTWATPKLSEDKVKQCVDSRLGGDYPPKAVAK-------------
AtPti1-8    273 RKPVDRTLPRGQQNLVTWATPKLSKDKVKQCVDARLLGEYPPKAVAK-------------
SlPti1      272 RKPVDHTLPRGQQSLVTWATPRLSEDKVKQCVDARLNTDYPPKAIAK-------------
AtCARK1     273 RKPVDHTLPRGQQSVVTWATPKLSEDKVKQCVDARLNGEYPPKAVAK-------------
AtPti1-11   272 RKPVDHTLPRGQQSLVTWATPKLSEDKVKQCVDARLLGEYPPKAVGK-------------

AtPti1-6    300 -------------------LAAVAALCVQYESNCRPKMSTVVKALQQLLIATGSIPQF--
AtPti1-1    321 -------------------LAAVAALCVQYEAEFRPNMSIVVKALQPLLKPPAAAPAPES
AtPti1-2    324 -------------------LAAVAALCVQYESEFRPNMSIVVKALQPLLKPPAPAPAPVP
AtMAZZA     366 -------------------LAAVAALCVQYESEFRPNMSIVVKALQPLLRSSTAAAVPVQ
AtCARK6     398 -------------------LAAVAALCVQYEAEFRPNMSIVVKALQPLLRSATAAAPPTP
AtMARIS     325 -------------------LAAVAALCVQYEADFRPNMTIVVKALQPLLNSKPAGPESTS
AtPti1-4    337 RFRLHSLFLTSSYGDDDSQLAAVAALCVQYEADFRPNMSIVVKALQPLLNARAVAPGEGV
AtPti1-9    320 -------------------LAAVAALCVQYEADFRPNMSIVVKALQPLLNARTGPAGEGA
AtPti1-8    320 -------------------LAAVSARCVHYDPDFRPDMSIVVKALQPLLNSSRSSPQTPH
SlPti1      319 -------------------MAAVAALCVQYEADFRPNMSIVVKALQPLLPRPVPS-----
AtCARK1     320 -------------------LAAVAALCVQYEADFRPNMSIVVKALQPLLNPPRSAPQTPH
AtPti1-11   319 -------------------LAAVAALCVQYEANFRPNMSIVVKALQPLLNPPRSAPQTPH

AtPti1-6        ----
AtPti1-1    362 ----
AtPti1-2    365 ES--
AtMAZZA     407 EA--
AtCARK6     439 QP--
AtMARIS     366 ----
AtPti1-4    397 H---
AtPti1-9    361 P---
AtPti1-8    361 WNPY
SlPti1          ----
AtCARK1     361 RNPY
AtPti1-11   360 RNPY
```
